# Rhythmic modulation of prediction errors: a possible role for the beta-range in speech processing

**DOI:** 10.1101/2022.03.28.486037

**Authors:** Sevada Hovsepyan, Itsaso Olasagasti, Anne-Lise Giraud

## Abstract

Natural speech perception requires processing the current acoustic input while keeping in mind the preceding one and predicting the next. This complex computational problem could be handled by a multi timescale hierarchical inferential process that coordinates information flow up and down the language hierarchy. While theta and low-gamma neural frequency scales are convincingly involved in bottom-up syllable-tracking and phoneme-level speech encoding, the beta rhythm is more loosely associated with top-down processes without being assigned yet a specific computational function. Here we tested the hypothesis that the beta rhythm drives the precision of states during the speech recognition hierarchical inference process. We used a predictive coding model that recognizes syllables *on-line* in natural sentences, in which the precision of prediction errors is rhythmically modulated, resulting in alternating bottom-up vs. top-down processing regimes. We show that recognition performance increases with the rate of precision updates, with an optimal efficacy in the beta range (around 20 Hz). The model further performs when prediction errors pertaining respectively to syllable timing and syllable identity oscillate in antiphase. These results suggest that online syllable recognition globally benefits from the alternation of bottom-up and top-down dominant regime at beta rate, and that the gain is stronger when different features are also analyzed in alternation. These results speak to a discontinuous account of inferential operations in speech processing.

## Introduction

A key challenge in speech processing is the ability to analyze what has just been said while processing what is being said and predicting what will follow, the so-called “now or never bottleneck” (Christiansen and Chater, 2016). This threefold challenge does not only require an appropriate neural architecture but also an efficient temporal orchestration of the neural event sequence involved, allowing through an inferential process for joint information intake, processing and prediction. This inferential process takes place in a left-hemispheric network (Rauschecker and Scott, 2009; Hamilton *et al.*, 2021; Zaccarella, Papitto and Friederici, 2021) where information flows up and down the hierarchy via feedforward and feedback connections and spreads at each stage via lateral connections (Friston, 2008; Friston and Kiebel, 2009; Cope *et al.*, 2017). Speech recognition results from the precise interplay between these feedforward, feedback and lateral streams during the multi-level inference (Davis and Johnsrude, 2007; Friston and Kiebel, 2009; Parr and Friston, 2018). Whether the inferential process involves continuous or discrete/alternating operations, and at which rate(s) they possibly occur is an essential piece of the puzzle.

Neural oscillations, as a proxy of rhythmic collective properties of neurons (Hauk, Giraud and Clarke, 2017; Meyer, Sun and Martin, 2019; Bree *et al.*, 2021), are directly involved in various aspects of speech processing (Zoefel, Archer-Boyd and Davis, 2018; Marchesotti *et al.*, 2020), including speech chunking at different granularity levels depending on their frequency (phrases, words, syllables, phonemic features) and information encoding depending on their cross-frequency interactions (Giraud and Poeppel, 2012; Hyafil *et al.*, 2015; Bonhage *et al.*, 2017; Ghitza, 2020; Proix *et al.*, 2022). Theta (4-7Hz) and low-gamma (25-35Hz) oscillations are related to bottom-up processes, notably the hierarchical encoding of phonemic information within syllables (Hyafil *et al.*, 2015; Mai, Minett and Wang, 2016; Lizarazu, Lallier and Molinaro, 2019). Delta (1-4Hz) and low-beta (14-21Hz) oscillations, which are also frequently observed in relation with speech processing, have a more endogenous origin. While delta is argued to play a role in syntactic parsing (Ding *et al.*, 2015, 2017), beta (15-30Hz) oscillations are associated with comprehension and top-down effects, without being related to specific linguistic units or language operations (Lewis and Bastiaansen, 2015; Pefkou *et al.*, 2017; Keitel, Gross and Kayser, 2018; Terporten *et al.*, 2018; Abbasi and Gross, 2020).

The notions of neural oscillations and hierarchical inference are likely intimately related to cognitive processes, notably in speech reception (Arnal and Giraud, 2012; Cope *et al.*, 2017; Abbasi and Gross, 2020; Donhauser and Baillet, 2020). Experimental studies and theoretical proposals suggest that information is generally transferred up and down the hierarchy using different frequency channels (Arnal and Giraud, 2012; Bastos *et al.*, 2012; Fontolan *et al.*, 2014; Chao *et al.*, 2018). Gamma oscillations (30-100Hz) are related to bottom-up information and prediction errors, i.e. the discrepancy between cognitive expectations and sensory signals (Lam *et al.*, 2016; Sedley *et al.*, 2016; Chao *et al.*, 2018), whereas beta oscillations (15-30Hz) are rather associated with top-down predictions and modulatory signals (Fontolan *et al.*, 2014; Arnal, Doelling and Poeppel, 2015; Chao *et al.*, 2018; Bastos *et al.*, 2020). The exact computational function of the latter, however, and their possible interplay with upgoing signals remains unclear (Bastos *et al.*, 2012; Fujioka *et al.*, 2012; Weiss and Mueller, 2012; Fries, 2015; Chang, Bosnyak and Trainor, 2018; Betti *et al.*, 2020).

Several hypotheses have nevertheless been formulated (Spitzer and Haegens, 2017; Miller, Lundqvist and Bastos, 2018; Betti *et al.*, 2020). Beta could work as an information channel conveying predictions down the processing hierarchy (Bastos *et al.*, 2015; Michalareas *et al.*, 2016), or, according to the predictive routing hypothesis, it could also prepare specific pathways by inhibiting neural populations that encode expected sensory signals, lowering the processing cost of novel information (Bastos *et al.*, 2020; Sherfey *et al.*, 2020). Not incompatibly, it might also reflect the delay for integrating bottom-up sensory signals and updating predictions (Arnal and Giraud, 2012). In the same vein, recent work suggests that beta oscillations could directly be related to the weighting of sensory prediction errors (Palmer *et al.*, 2019).

Following-up on this, we used computational modeling to address the possible function of beta oscillations in the rhythmic weighting of prediction error in the context of speech processing. We built on a previous model that uses theta (~5Hz) / gamma (~40Hz) oscillation coupling in a predictive coding framework to achieve natural speech parsing and *on-line* syllable identification in continuous natural speech (Hovsepyan, Olasagasti and Giraud, 2020). In the new model, *Precoss-β*, we explore how alternating top-down and bottom-up information streams via the rhythmic weighting of prediction errors affects the inference process.

Modulating prediction error precisions (PEP) within a frequency range spanning from 2 Hz to 60 Hz, for both syllable identity and timing, we found that *Precoss-β* outperformed its previous version with non-modulated prediction errors, and was most efficient when precisions were modulated at the beta range (20-30Hz). These results show that the low-beta rhythm supports online speech recognition by controlling the alternation of a bottom-up versus top-down dominant mode during the inference process. The observed benefit reflects that the model can flexibly pick up unexpected input while remaining both sensitive to bottom-up information and reliable in terms of predictions, hence achieving the triple challenge of speech processing.

## Results

### *Precoss-β* architecture and oscillating precisions

*Precoss-β* was built by including oscillating state-dependent precisions in a previous generative model version (*Precoss*) (Hovsepyan, Olasagasti and Giraud, 2020). The model input consists of a speech reduced auditory spectrogram (Chi, Ru and Shamma, 2005) and of its slow amplitude modulations (Hyafil *et al.*, 2015), both extracted English sentences of the TIMIT database (Garofolo *et al.*, 1993) (see Hovespyan et al. 2020 (Hovsepyan, Olasagasti and Giraud, 2020) for details about speech input generation).

In *Precoss-β*, the activation of the appropriate syllable unit generates the corresponding auditory spectrogram with a flexible duration determined by eight gamma units (Figure 1). Syllable and gamma units represent syllable identity and timing within the syllable, respectively. Together with the other model elements, they are used to deploy predictions about the input acoustic spectrogram. The ongoing mismatch between predicted and actual auditory spectrograms and slow amplitude modulations drives the inference process across the model hierarchy and leads to updating syllable and gamma units (as well as all other variables in the model) such that predictions best match the input.

**Figure 1:**
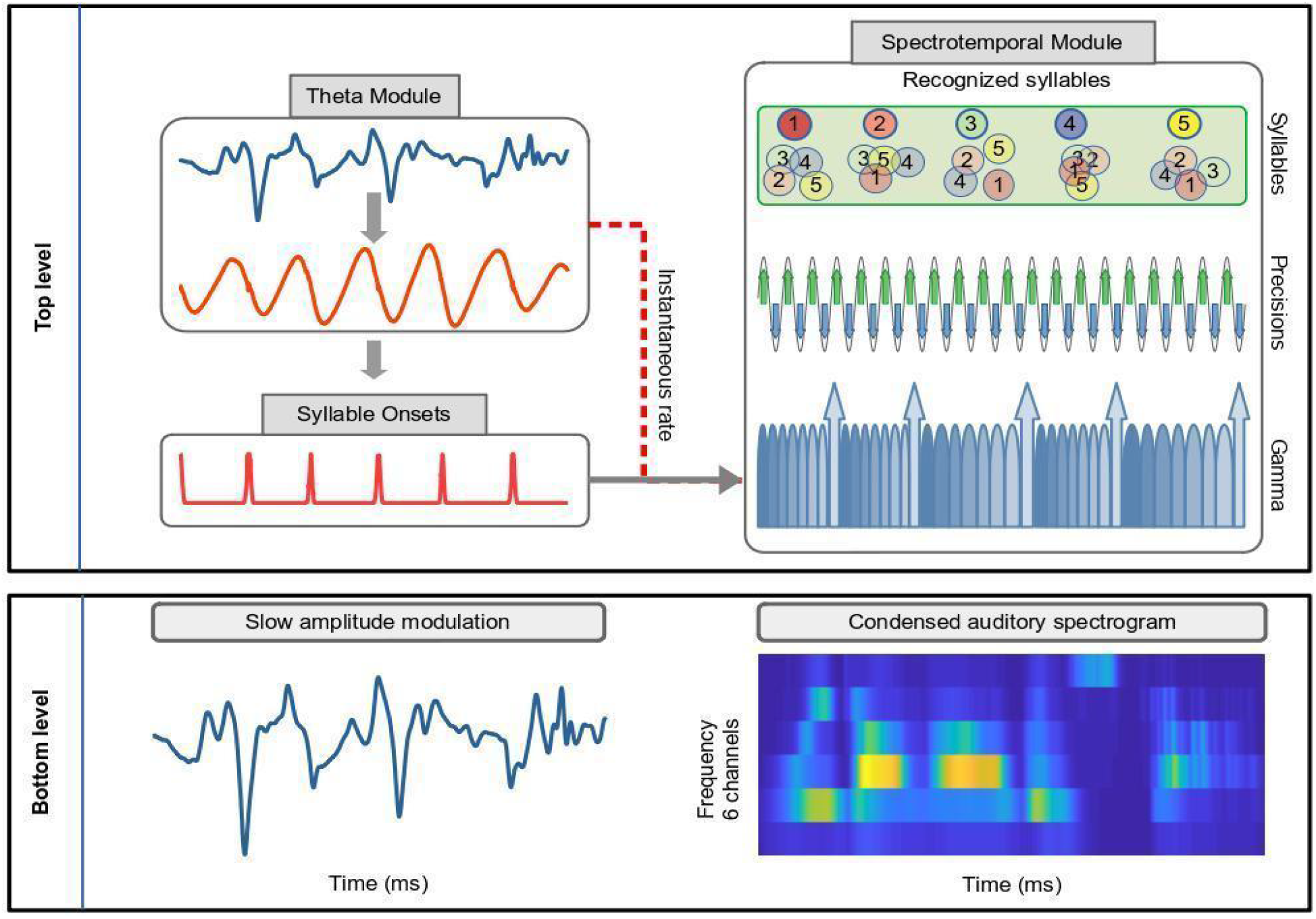
A generative model for on-line syllable recognition with rhythmic state-dependent precisions. The bottom panel represents the input to the model. As in the original model (*Precoss*), the input consists of the speech slow amplitude modulation (on the left) and auditory spectrogram (on the right). On the top level, the theta module tracks the slow amplitude modulation in the input and feeds it to a theta oscillator. Theta-triggers, associated with a pre-defined phase, signal the estimated syllable onsets which reset the sequence of gamma units in the Spectrotemporal module. Additionally, the instantaneous frequency of the theta oscillator sets the preferred rate of the gamma sequence. Together, gamma and syllable units generate the auditory spectrogram in the input. Accumulated evidence about each syllable in the sentence, represented by syllable units, is reset by the last gamma unit (upwards arrow). Precision units change the precision of syllable and gamma units, modulating the influence of the corresponding prediction errors on the hidden states’ dynamics. Depending on the phase of precision units (green and blue arrows), either syllable or gamma units get higher precision (*Precoss-β* (C)).

As our goal is to assess how rhythmic fluctuations of internal expectation vs. bottom-up prediction errors drive the model updates with respect to syllable identity (syllable units) and timing (gamma units), and affect performance, we introduced specific units that control the precision of syllable and/or gamma units (variants A, B and C in Figure 2). These precision units effectively modulate the relative strength of internal predictions based on previous time points and bottom-up prediction errors in the updates of syllable and/or gamma units.

**Figure 2:**
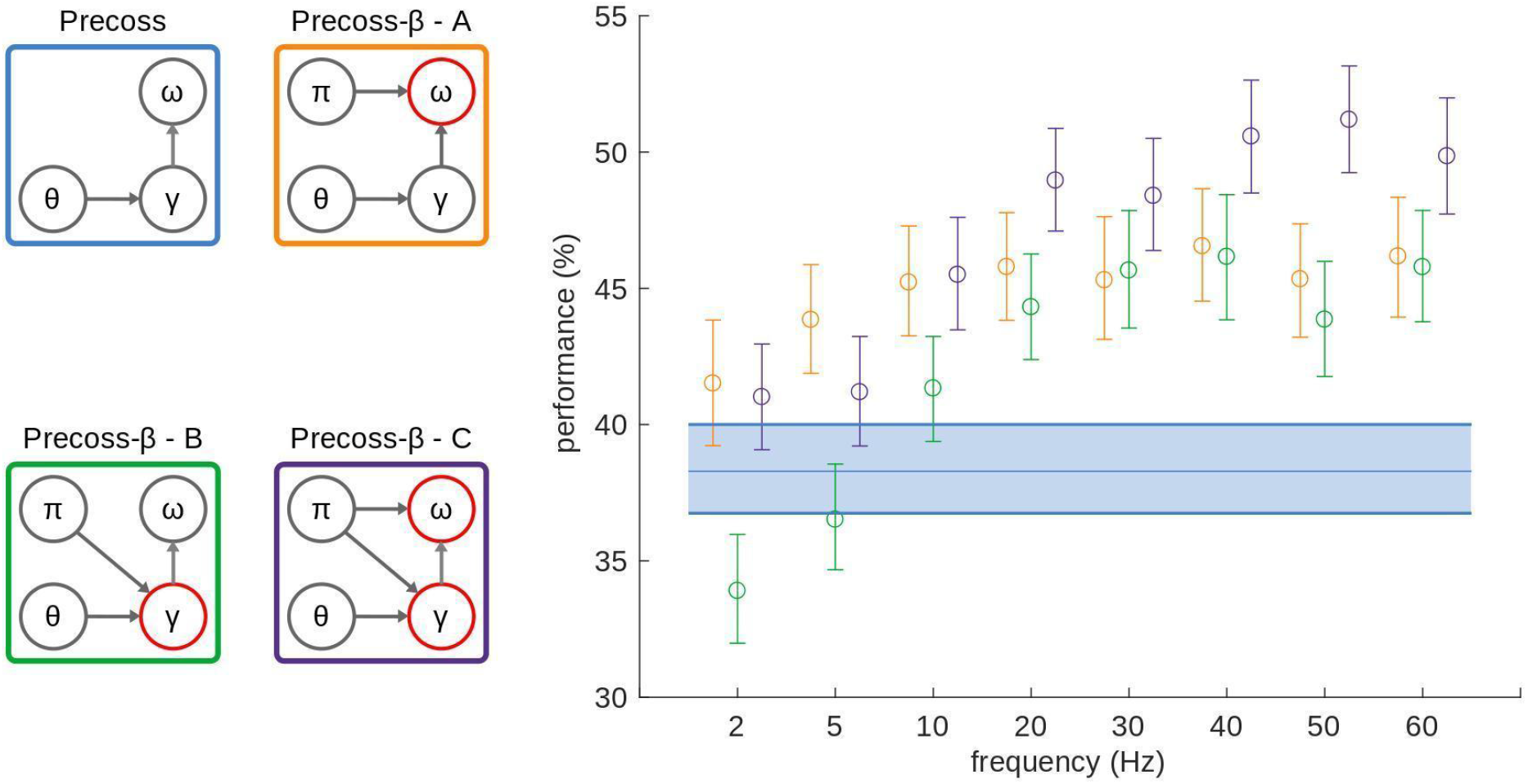
Model performance based on the overlap measure. We tested the online syllable recognition accuracy of the model based on simulation results on 220 sentences (giving a total of about 3000 syllables). Accuracy was evaluated based on the overlap of the recognized syllable sequence and the input sentence. Data for each model variant is represented by the colour of the outlines on the left panel. The figure shows the mean performance and 95% confidence interval for each frequency value of precision units. Diagrams on the left indicate the main functional groups of the model: θ corresponds to the theta-module, γ and ω to syllable and gamma units respectively. Arrows indicate connections between functional groups (θ → γ represents rate and onset information from theta module to gamma units, whereas γ → ω indicates the reset of accumulated evidence by the last gamma unit). π represents precision units, and the arrows originating from it indicate which functional groups they control.

The model performance was assessed via the output of syllable units, which summarizes the model estimate about the syllable boundaries and identity in the speech input. Performance measures are based on comparing the estimated syllable sequence with the actual one in the input.

### Model variants and performance

To assess the effect of modulating top-down and bottom-up information streams, we compared the performance of *Precoss* (stationary precisions) and *Precoss-β* (oscillating precisions) in their ability to parse and recognize syllables from natural spoken sentences. Whatever the model, the input is a full sentence without explicit syllable boundaries, and the model parses it into discrete units and identifies the sequence of activated syllables.

Since predictions about the auditory spectrogram (the input) are generated in concert by syllable units that recognize the overall spectrotemporal pattern, and gamma units that specify the position of the acoustic segment within the overall pattern, the discrepancy between the predicted and actual input can in principle be solved by updating both the estimate of *where* we are in the pattern (gamma units) and the pattern identity (*what* - syllable units).

We, therefore, run simulations varying the frequency at which precision units modulate syllable and gamma units. We compared model variants (Figure 2, left panel) where precisions drive: causal syllable units alone (*Precoss-β* (A)), causal gamma units alone (*Precoss-β* (B)) or both in anti-phase (*Precoss-β* (C)). In the latter case, anti-phase refers to the fact that when syllable units are in a high precision state, gamma units are in a low precision one, and vice versa. We also considered the case where both causal gamma and syllable units are in phase (*Precoss-β* (C’), Figure 5). The original model with stationary precisions provides baseline performance. The simulations were run on the same set of 220 natural sentences.

We posit that modulating the relative strength of internal expectation and bottom-up information in a rhythmic fashion is expected to improve performance as it alternatively sensitizes the model to internal knowledge vs. external evidence, which, given the predictive nature of speech, should be an optimal processing strategy. We also expected the model performance to depend on the prediction error precision rate, peaking at a frequency that will depend on the two model intrinsic rhythms (~ 200 ms for syllable units, ~ 25 ms for gamma units).

In *Precoss-β* (A) PEP are only modulated in the syllable units, which act as evidence accumulators for each syllable in the input sentence. Therefore, to benefit from the alternation between top-down and bottom-up information flows on the inference process, there should be at least one full PEP cycle per syllable. As the mean syllable duration in our dataset is around 200ms, we expect the preferred PEP modulation frequency to lie within the theta range ~ 5 Hz.

Similarly, in *Precoss-β* (B) PEP are only modulated within the gamma units. Those units are responsible for deploying spectrotemporal predictions at the right time and in the correct order. They operate at gamma scale (40 Hz, at rest). With the same logic as for variant A, we thus expect a positive effect on alternation to require a PEP modulation frequency within the gamma range.

Finally, in *Precoss-β* (C) PEP are modulated in both syllable and gamma functional groups. As information syllable identity fluctuates at the theta range and information about timing fluctuates at the higher gamma range, we expect the optimal common PEP frequency to lie somewhere between 5 Hz and 40 Hz.

Figure 2 shows the performance of variants A, B, and C together with that of the original *Precoss* with stationary precisions. To quantify syllable recognition performance we compared the model output and input. The performance metric takes into account both the order and duration of the syllables and varies between (0-100%) (for details about this measure see the original paper and supplementary Figure 1 (Hovsepyan, Olasagasti and Giraud, 2020)). For almost all conditions, *Precoss-β* (oscillating precisions) significantly (Supplementary Tables 1-3) outperformed *Precoss*. That is, the rhythmic alternation of internal expectations and bottom-up influence on the inference process improves online syllable recognition from natural sentences.

The orange dots and ranges represent the mean performance and 95% confidence intervals for *Precoss-β* (A) obtained by bootstrapping with 10000 reps. For all tested PEP modulation frequencies, *Precoss-β* (A) performed better (Wilcoxon signed rank test, Z=4.78, p=1.784e-6, at 5 Hz) than *Precoss* with stationary precisions (blue line). The difference was statistically significant (p<0.05) for all frequency values (Supplementary Table 1). However, no optimal frequency arose; performance reached a plateau at 5Hz and fluctuations beyond 5Hz were not statistically significant (Supplementary Table 4).

Simulation results for *Precoss-β* (B) are presented in green. Interestingly *Precoss-β* with oscillating precisions performed lower than *Precoss* with stationary precisions for low modulation frequencies (Wilcoxon signed rank test, Z=-3.653, p=2.586e-4 at 2 Hz) and higher for modulations >10 Hz does (Wilcoxon signed rank test, Z=5.55, p=2.819e-8 at 20 Hz) (Supplementary Table 2). Although performance peaked in the gamma range (Wilcoxon signed rank test, Z=6.3, p=2.81e-10 at around 40Hz), pairwise comparisons were not statistically significant for frequencies equal or greater than 20 Hz, indicating a knee point at this frequency (Supplementary Table 5).

Finally, *Precoss-β* (C), which controls precisions of both syllable and gamma units, outperformed *Precoss* for all frequency values (Supplementary Table 3). Here again, we do not see a preferred frequency for the best model performance, instead, performance increases with frequency and reaches a plateau at around 20 Hz (Wilcoxon signed rank test, Z=9.21, p=3.205e-20). Interestingly, while for lower frequencies *Precoss-β* (A) and (C) perform more or less similarly, for frequencies higher than 20 Hz, *Precoss-β* (C) outperforms the other model variants (Supplementary table 7). As was the case for *Precoss-β* (B), pairwise comparisons of model performance for different frequencies higher or equal to 20 Hz, were not statistically significant (Supplementary Table 6).

### Bayesian information criterion

The model performance based on the overlap measure evaluates the model’s ability to correctly infer syllables’ identity and duration in the input sentence. However, neither the model complexity - e.g. Precoss-β vs. Precoss, nor the uncertainty of the model’s estimates is taken into account in performance. We thus also calculated the Bayesian Information Criterion (BIC) (Schwarz, 1978) for each model variant and each PEP modulation frequency.

BIC was evaluated from the dynamics of the syllable units (not averaged within gamma-defined windows) as well as the model estimated conditional precisions at each time point (Hovsepyan, Olasagasti and Giraud, 2020). Thus BIC value represents how the syllable units’ accuracy and related model-estimated uncertainty change with the PEP modulation frequency.

Results are presented in Figure 3. Despite higher performance for *Precoss-β* than *Precoss* when based on the overlap measure represented in Figure 2, BIC penalizes variants (A) and (B) relative to *Precoss* at all frequencies. However, *Precoss-β* (C), which controls the precision of both syllable and gamma units, consistently displays a higher BIC value than *Precoss* (despite having more parameters) and other equally complex *Precoss-β* variants. This suggests that controlling the precision of both functional groups does not only result in higher syllable recognition but also in higher confidence in the inference. Interestingly, BIC peaks at 10 Hz, closely followed by 5 Hz and 20 Hz. This range aligns well with results from electrophysiological studies showing the alpha-beta range in speech tasks explicitly involving inferential processes (Mai, Minett and Wang, 2016; Sedley *et al.*, 2016; Pefkou *et al.*, 2017)

**Figure 3:**
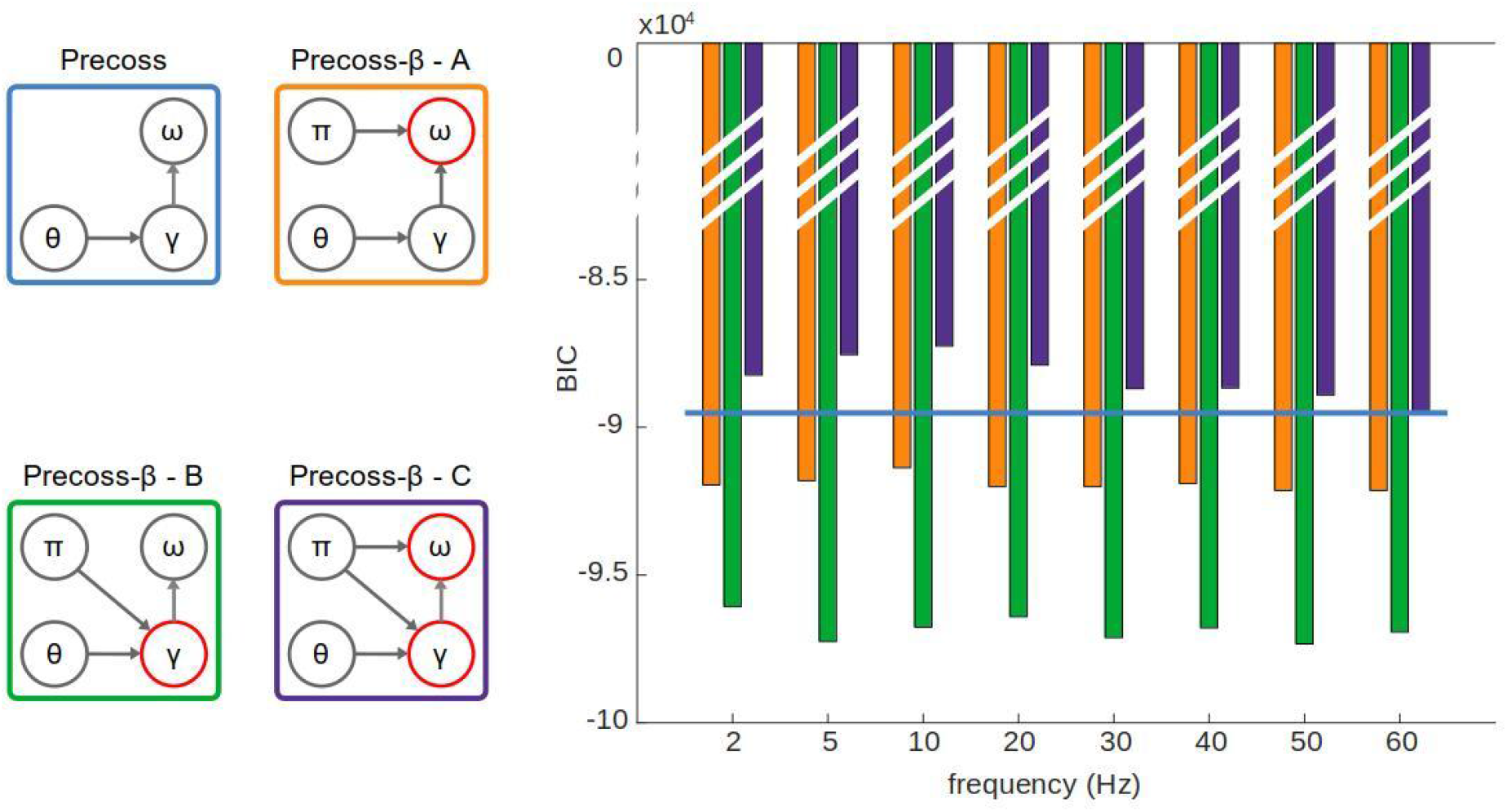
Bayesian information criterion. The left panels represent the Precoss-β variants with oscillating precisions and *Precoss* as in Figure 2. *Precoss-β* (C) has the highest BIC values among all models at precision frequency 5-20 Hz. These results suggest that the model is the most confident about its inference when the influence of the top-down and bottom-up information flows on the model’s dynamics alternates within this specific frequency range.

### Modulation of bottom-up information

Another important factor in evaluating the model is the informativeness about syllable identity propagated up in the hierarchy via bottom-up prediction errors, and how it might be affected by PEP modulation frequency. We hence quantified the informativeness of bottom-up prediction errors by taking into account whether prediction errors signaling a syllable in the input arrive when the syllable is already in a high activity state (low informativeness) or still in a low one (high informativeness).

The results are presented in Figure 4. Frequency significantly affected the informativeness measures (Friedman test, χ^2^ = 269.85, p = 1.635e-54), which was was significantly higher at 20 Hz than at all other frequencies except 30 and 40 Hz (Bonferroni corrected post-hoc pairwise comparisons, see Supplementary Table 11 for details). These results confirm the beta range as an efficient modulation frequency for alternating the influence of top-down and bottom-up information.

**Figure 4:**
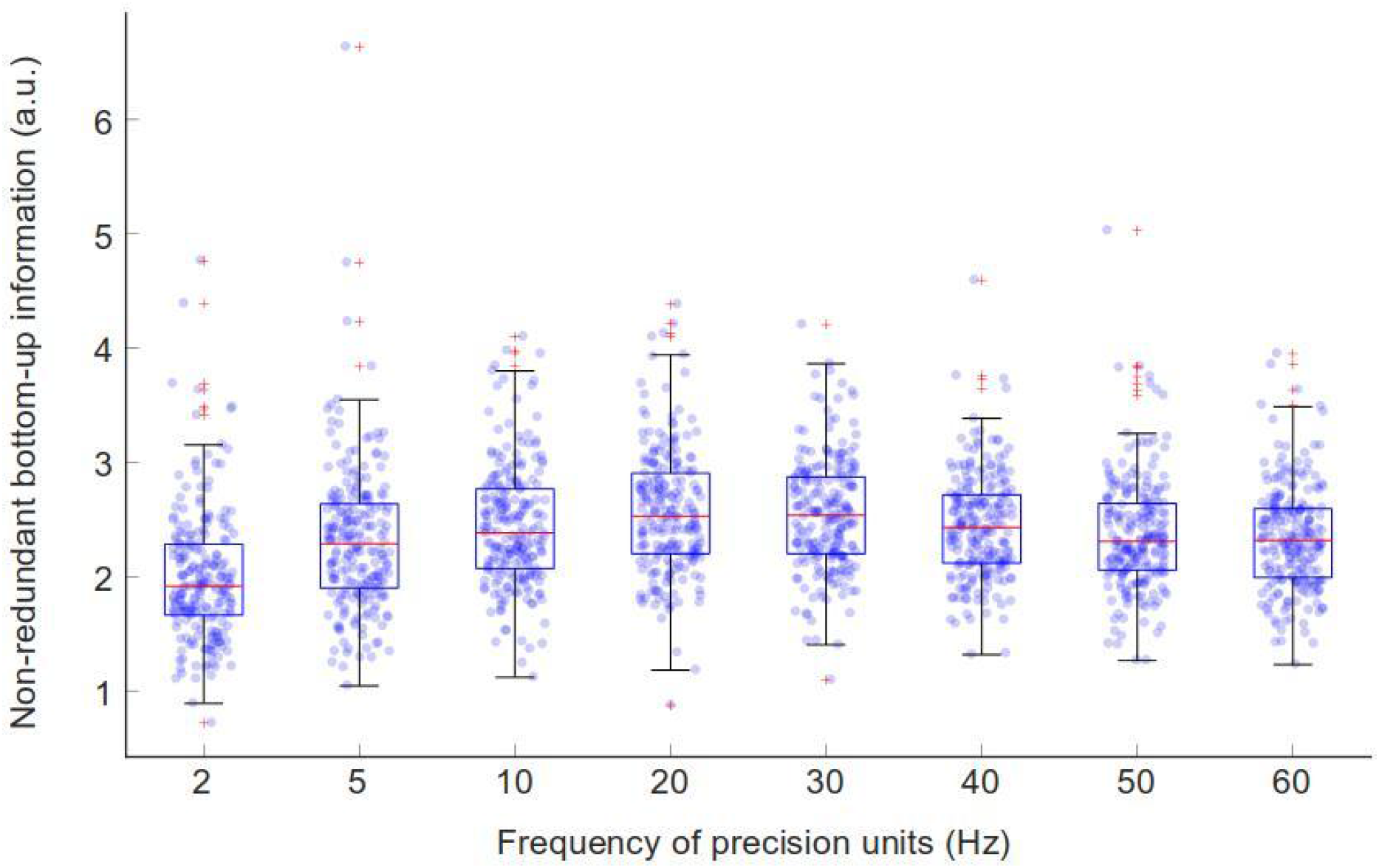
Non-redundancy of bottom-up information - *Precoss-β* (C). We evaluated how the modulation frequency of PEP affects the amount of informative (non-redundant) bottom-up signal. Friedman test indicated that the modulation frequency affects the amount of informative signal propagated up in the model-hierarchy. Pairwise comparisons were performed for each frequency pair (Supplementary Table 11). Bonferroni procedure was used to control for multiple comparisons. The measure for non-redundant bottom-up signal peaked at 20 Hz, with statistically significant (p<0.05) differences from all other frequencies except 30 and 40 Hz. Each point on the scatter plot represents the measurement value for each sentence for the corresponding PEP frequency. The central-red mark of the box plots corresponds to the median, whereas bottom and top edges represent 25th and 75th percentiles, respectively. Red crosses indicate outliers, whereas whiskers extend to the highest and lowest performance values that are not considered outliers.

### Effect of PEP modulation phase

Among the three model variants, the best performance (highest number of recognized syllables with the least uncertainty) is obtained for the one where PEP are modulated in both syllable and gamma units. By construction, this model variant (C) controls the precisions of syllable and gamma units in opposite directions; whenever the precision of syllable units increases the precision of gamma units decreases and vice versa. This choice was based on the idea that syllable units and gamma units can take turns in *absorbing* prediction errors, making it easier for the model to find the right estimates.

To address how this *a priori* choice affected performance, we also run the model with precisions of gamma and syllable units oscillating in-phase (same-phase condition, Figure 5, red diagram on the left). The model with anti-phase condition outperformed the model with same-phase conditions at all frequencies (Figure 5, right panel, Supplementary table 8). This finding shows that the model performs best when bottom-up prediction errors are preferentially minimized in alternation by syllable and gamma units, when syllable identity and timing features are analyzed via antagonistic streams.

**Figure 5:**
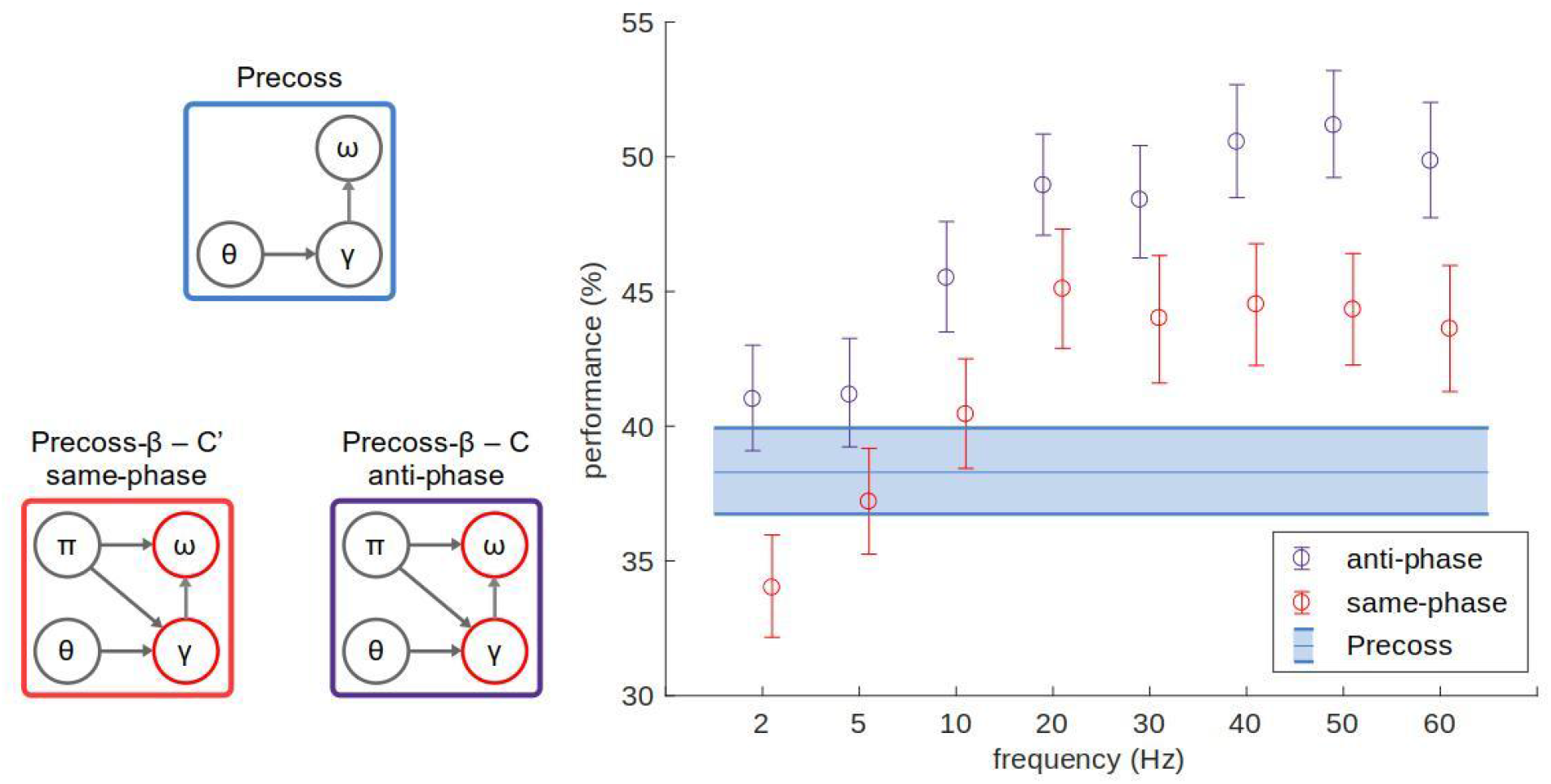
Effect of the oscillating PEP phase on model performance. *Precoss-β* (C) controls the precision of both syllable and gamma units so that the high precision state of the syllable units coincides with the low precision state for gamma units (anti-phase condition, purple). Here we tested whether the performance depends on the phase lag for the precisions of the syllable or gamma unit. Thus, we also tested *Precoss-β* (C’) when syllable and gamma units reach a high precision state simultaneously (same-phase condition, red). The mean values (based on the overlap measure, supplementary figure 1) and corresponding 95% confidence intervals are presented in the figure. Results suggest that for all frequencies, the anti-phase condition leads to better model performance. These results indicate that high-versus low-precision states should oscillate in different directions for different functional (*what* vs. *when*) groups.

## Discussion

The goal of this study was to explore the possible role of cortical beta oscillations in speech processing from a theoretical perspective, where the brain deploys predictions through top-down connections and refines them based on bottom-up prediction errors (Rao and Ballard, 1999; Friston and Kiebel, 2009; Shipp, 2016). Here, we conjectured that beta oscillations might reflect the alternation of bottom-up versus top-down control in the brain’s inference process, and tested this hypothesis by introducing precisions that oscillated in time within feature-specific functional groups (syllable recognition and timing units) and comparing performance across frequencies with a baseline/control model with stationary precisions. We found that performance improved with the PEP oscillation rate, and reached a plateau, with a knee point that depends on the model variant (precision of which functional group is controlled).

### The added value of rhythmic prediction error precisions (PEP)

The model encompasses two distinct functional groups operating in two distinct regimes: when the causal states of one group (syllable and/or gamma units) are in the low precision phase of the oscillation, they are both less strongly receptive to the internal expectations encoded by the hidden states and more strongly influenced by the bottom-up input carrying prediction errors from the periphery. As a result, each functional group is periodically in an optimal position to respond to bottom-up information without being constrained by internal expectations. And vice versa in the high-precision phase, where causal states are preferentially coupled to hidden states encoding internal expectations and more loosely to bottom-up input. The high-precision phase is therefore ideal to incorporate updates from the preceding low-precision phase into the internal hidden states. Thanks to oscillating PEPs, the model is rhythmically alternating between an information gathering and an information consolidation regime. The newly consolidated information leads to updated predictions, which in the next cycle are again compared with the input leading to updates in causal states, and to a new round of consolidation. That *Precoss-β* outperformed *Precoss* for almost all PEP frequencies indicates that rhythmic alternation of top-down and bottom-up streams during the inference process improves online syllable recognition. An important issue is therefore whether there is an optimal oscillating PEP rate in speech processing.

### Low-beta as an optimal range for rhythmic PEP

The different variants of *Precoss-β* were assessed based on three different metrics. One that assesses syllable accuracy and duration (Figure 2), another that additionally takes into account the number of parameters of the model and the confidence in its output (Figure 3), and one that quantifies the amount of non-redundant bottom-up information about syllable identity (Figure 4).

Our initial hypothesis was that performance would take a bell shape as a function of frequency, indicating an optimal PEP oscillation regime, depending on whether syllable vs. gamma units or both were modulated. Our reasoning was that when PEP rhythm is lower than the intrinsic rhythm of syllable and gamma units, there is less than one full PEP cycle per information cycle. Conversely, for frequencies higher than the average rate of syllable and/or gamma units, the alternation of high and low precision states becomes too fast for the model to grasp relevant bottom-up information, leading to a weaker performance.

Yet, Figure 2 shows that performance for all model variants (as assessed by syllable accuracy and duration) reaches a plateau rather than showing a peak frequency. Only the knee point of the plateau differed from variant to variant: 5 Hz for *Precoss-β* (A), which roughly corresponds to the natural syllabic rhythm, and 20 Hz for *Precoss-β* (B), a relatively sensible result given that gamma units are designed as a stable heteroclinic channel where activity within neighboring units can overlap in time.

For *Precoss-β* (C), while we also observed the knee point at 20 Hz, the performance attained was higher than for *Precoss-β* (A, B), indicating an additive benefit of controlling PEP in both syllable and gamma units. This additive effect occurs when the modulation of syllable and gamma units are in anti-phase, i.e. when one functional group is in a high precision state while the other is in a low one. In the anti-phase condition, only one functional group at a time (the one in the low-precision phase) can grasp changes in the input, while the other incorporates information from the causal state into the dynamics. This alternation regime reduces the search space compared to the variant where the model tries to optimize syllable and gamma units simultaneously (Figure 5).

Theoretically, the appropriate rhythm to control precisions within early speech processing stages should hence be both slow enough to span across processing stages (and modules) and fast enough to achieve an optimal balance between input sensitivity and prediction updating. The beta rhythm, as an intermediate range between theta and gamma, might hence be ideally suited for both purposes. The model accordingly reaches a plateau at 20 Hz (low-beta range). Even though in terms of raw performance, higher PEP frequency might result in better performance, lower beta-range frequencies might be more suited in a real hierarchical structure. Accordingly, beta oscillations are considered to be a good channel for long-range communication (Kopell *et al.*, 2000; Engel and Fries, 2010; Spitzer and Haegens, 2017; Betti *et al.*, 2020). Beta oscillations that originate in higher levels of the cortical hierarchy could update precisions via a cascade running through the whole hierarchy down to the sensory areas.

Although assessing the model performance did not allow us to identify an optimal rhythm, but only the lower-bound of an optimal PEP frequency range, the Bayesian Information Criterion (BIC, Figure 3), which took into account both confidence and the moment-to-moment dynamics of syllable units, showed a peak between 5-20 Hz for the best model variant, an interval that includes the cortical low-beta range (15-20 Hz). In addition, the estimate of bottom-up informativeness peaked at 20 Hz (Figure 4). These results indicate that the low-beta range (~20 Hz) fulfills three factors for successful inference: syllable recognition accuracy (Figure 2), inference confidence (Figure 3) and non-redundancy of bottom-up information about syllable identity.

### Rhythmic PEP and precision theories

Sensorimotor beta has been linked to precisions before (Tzagarakis *et al.*, 2010; Tan, Wade and Brown, 2016). Sensorimotor beta activity is argued to reflect the integration of the sensory signal uncertainty with the uncertainty of the internal model about prediction errors (Palmer *et al.*, 2019). Here, we confirm the implication of the beta rhythm in this process, and go further in showing 1) that the rhythmic modulation of precisions changes the relative weight of bottom-up vs. top-down information during the inference process, and 2) that this is beneficial in an eminently dynamic task such as online syllable recognition. In other words, while precision (via e.g. synaptic gain) is important to assign uncertainty about the input throughout the hierarchy, there is an added benefit when it oscillates. Given bottom-up processes in the gamma and theta ranges, beta oscillations provide an optimal *timescale* to update precisions.

In sum, the role of beta oscillation (or more generally the notion of oscillating precisions) is to rhythmically modulate the relative influence of top-down and bottom-up information flows on the fly during a multi-level inference process, here hierarchical speech processing. Beta oscillations would hence not only act as an information channel (Bastos *et al.*, 2015; Michalareas *et al.*, 2016), but their effect on precisions would further modulate the strength of the top-down information flow.

### Rhythmic PEP and Predictive Routing

The rhythmic precision hypothesis is in line with studies suggesting rhythmic attentional sampling (Buschman and Miller, 2010; VanRullen, 2016, 2018; Fiebelkorn and Kastner, 2019). The *good* and *bad* phases associated with attentional sampling are conceptually similar to high and low precision states in the model. When bottom-up prediction errors have low precision, their contribution to the model dynamics decreases. This is similar to forming internal expectations while periodically scanning the sensory signal for something new or unexpected. Low precision phases provide windows of opportunities to detect new syllables in the input. In the absence of a new syllable, there is no substantial prediction error and the current syllable unit remains the most active one. Conversely, a new syllable triggers prediction error which will, at the next precision increase phase, switch the corresponding hidden state to its active form.

This scenario works when there are already internal expectations formed about the sensory signal. For example, when subjects listened to short stories, beta activity built up when more context became available (Pefkou *et al.*, 2017). As the current model does not include higher hierarchical stages (word, phrase levels) it implicitly assumes that expectations are already formed and that there is ongoing beta activity. This assumption is sufficient for the demonstration that oscillating precisions are helpful for online syllable recognition. However, in the brain, beta activity is most probably not always on, but rather appears as bursts of transient activity when top-down predictions are possible. Bastos and colleagues (2020) introduced *predictive routing* as an implementation of hierarchical processing during visual perception (Bastos *et al.*, 2020). Predictive routing assumes that alpha/beta bands prepare the pathways to process the predicted input by inhibiting bottom-up (non predicted) sensory information communicated at the gamma scale. Electrophysiological recordings showed enhanced alpha (8-14Hz) and beta (15-30Hz) activity for predictable stimuli, and gamma activity (40-90Hz) for unpredictable ones, especially in the lower layers of the hierarchy (Bastos *et al.*, 2020). These results may also be explained by beta activity controlling precisions; when the stimulus is predictable and internal expectations are formed, beta activity originating from higher cortical areas modulates precisions throughout the whole hierarchy, explaining more alpha/beta power across the hierarchy for predictable signals. For unpredictable stimuli, there are no internal expectations and no need for an alternated contribution of top-down and bottom-up streams. In this case, the system takes in sensory information with more bottom-up activity communicated by gamma oscillations. The predictive routing framework can in our opinion comfortably accommodate the notion that beta oscillations control state precisions, and mediate the contribution of top-down and bottom-up information during the hierarchical (inferential) perception process.

That our model performance was better when the precision of different functional groups of the model oscillates in opposite directions, speaks to an oscillatory regime spanning across various areas encoding different features. In our case, two distinct functional groups are responsible for predicting syllable identity and timing, mimicking the parallel implementation of what and when/where features in separate processing streams. We found better performance when the model controlled the PEP of these two functional groups in opposite directions, revealing higher computational efficiency when bottom-up sensory information is collected within one functional group, while the accumulated evidence is integrated into the hidden states in the other one. By reflecting rhythmic PEP modulation, beta activity may genuinely *represent* the beta rhythm that carries top-down information. How such a functional theory could be implemented at the biophysics levels remains to be established, but is not incompatible with models of beta ryhyhtm generation (Kopell *et al.*, 2000; Roopun *et al.*, 2008).

## Conclusion

This computational study suggests a new functional role of cortical oscillations during hierarchical syllable recognition from natural sentences. First, we show that online syllable recognition benefits from oscillating precisions that rhythmically modulate the contribution of top-down and bottom-up streams during the speech perception process. The performance gain is most tangible when one functional group integrates bottom-up information, while the other ignores it and only maintains internal expectations. The best performance is achieved when the model controls precisions across functional groups in the 20 Hz range, a range where we found the highest number of recognized syllables, higher confidence and less redundancy in bottom-up information about syllables. Oscillating PEP allows the model to reactively detect changes in the input, while also being able to maintain internal expectations, hence offering a new mechanistic role for the beta range in speech processing.

## Methods

### Speech input and syllabification

We have used the same set of 220 sentences from the TIMIT dataset (Garofolo *et al.*, 1993) that we used in (Hovsepyan, Olasagasti and Giraud, 2020) for the simulations of the new model - Precoss-β. Briefly, for each sentence, a 6-channel reduced auditory spectrogram was calculated with a biologically plausible model of the auditory periphery (Chi, Ru and Shamma, 2005). Additionally, slow amplitude modulation of the sentence waveform was calculated following procedures described in Hyafil and colleagues (Hyafil and Cernak, 2015; Hyafil *et al.*, 2015).

Syllable boundaries in the input sentences were defined with the Tsylb2 (Fisher, 1996) programme based on the phonemic transcriptions provided in the TIMIT database (Garofolo *et al.*, 1993). The programme estimates syllable boundaries based on English grammar rules, using phoneme annotations from TIMIT. Finally, syllable spectrotemporal patterns are calculated and stored in 6×8 matrices (6 frequency channels x 8 gamma units), where each row corresponds to the average value of the corresponding frequency bands within 8 binned temporal windows (assigned to specific gamma unit). For a detailed description of input construction and syllabification, please see the Methods section in (Hovsepyan, Olasagasti and Giraud, 2020).

### Generative model and *Precoss-β*

We use predictive coding to construct a model for parsing and recognizing syllables from natural English sentences. *Precoss-β* has the same hidden and causal states as in the original Precoss (Hovsepyan, Olasagasti and Giraud, 2020), but is defined with two additional hidden states at the top-level. These represent the harmonic oscillator that controls the precision of syllable and/or gamma units:

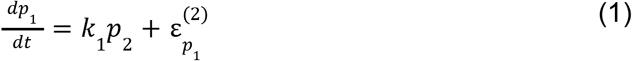

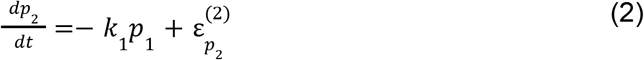

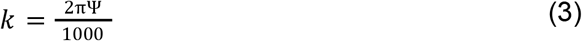

Equations 1 and 2 correspond to the oscillating precisions, Ψ in equation 3 corresponds to the modulation frequency of prediction error precisions in Hz and 1000 is the sampling rate. We have tested each Precoss-β variant for different values of the modulation frequency Ψ ranging from 2 Hz up to 60 Hz. Table 1 contains precisions for new hidden states and oscillating causal states for each model variant.

**Table 1:**
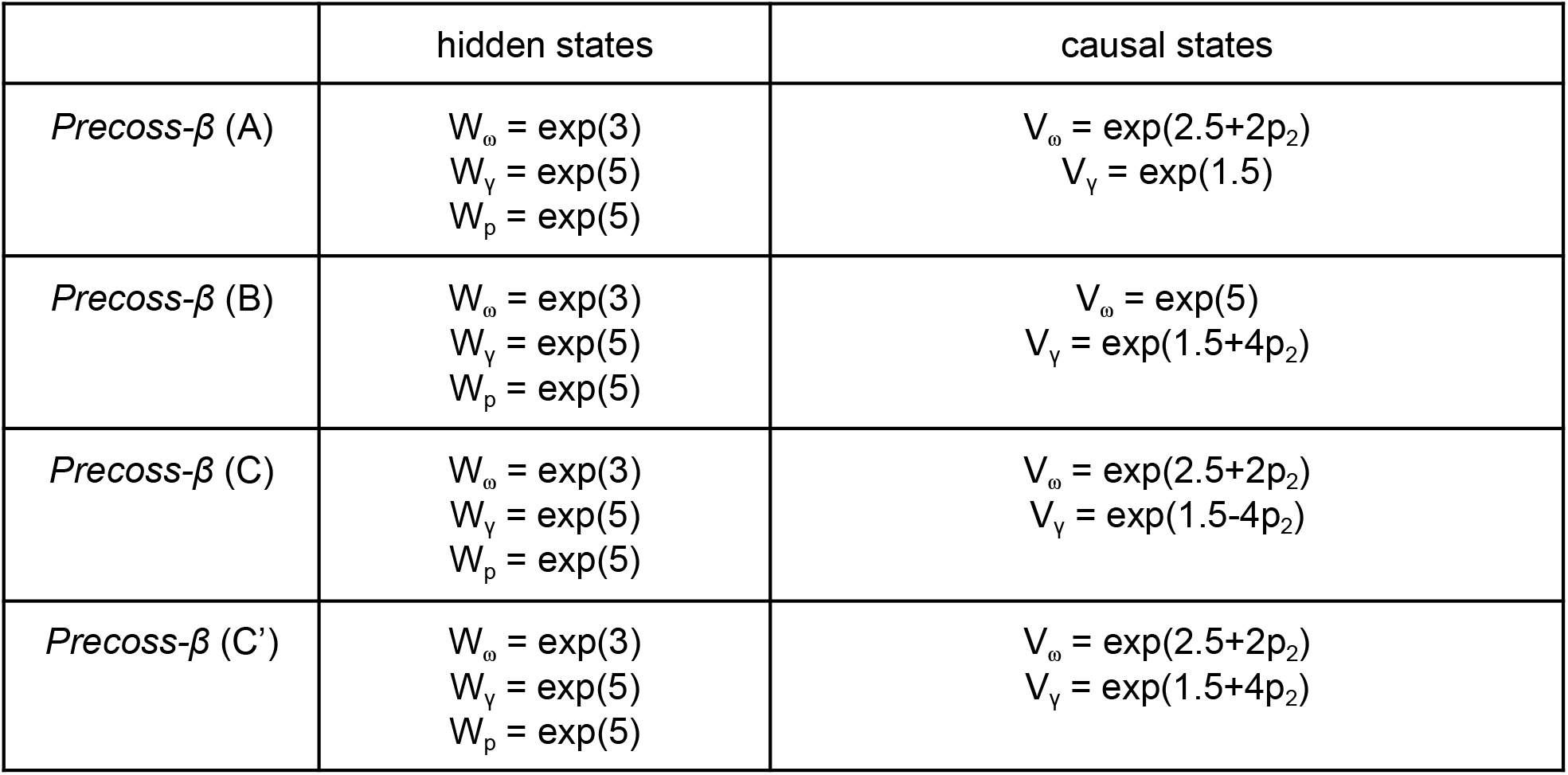
Precisions of syllable, gamma units and hidden states of the oscillating precisions. The left column represents stationary precisions for syllable and gamma units W_ω_ and W_γ_ respectively, and for the new hidden states that generate oscillating precisions - W_p_. The right column represents the precision of causal states for each variant. Depending on the Precoss-β variant either syllable (A) or gamma (B) units have oscillating precision. Meanwhile, for variants C and C’, both syllable and gamma units have oscillating precisions, with the difference that for variant C they oscillate in opposite phases, while for C’ in the same phase.

|  | hidden states | causal states |
| --- | --- | --- |
| *Precoss-β* (A) | $W_{\omega} = \exp(3)$<br>$W_{\gamma} = \exp(5)$<br>$W_p = \exp(5)$ | $V_{\omega} = \exp(2.5+2p_2)$<br>$V_{\gamma} = \exp(1.5)$ |
| *Precoss-β* (B) | $W_{\omega} = \exp(3)$<br>$W_{\gamma} = \exp(5)$<br>$W_p = \exp(5)$ | $V_{\omega} = \exp(5)$<br>$V_{\gamma} = \exp(1.5+4p_2)$ |
| *Precoss-β* (C) | $W_{\omega} = \exp(3)$<br>$W_{\gamma} = \exp(5)$<br>$W_p = \exp(5)$ | $V_{\omega} = \exp(2.5+2p_2)$<br>$V_{\gamma} = \exp(1.5-4p_2)$ |
| *Precoss-β* (C') | $W_{\omega} = \exp(3)$<br>$W_{\gamma} = \exp(5)$<br>$W_p = \exp(5)$ | $V_{\omega} = \exp(2.5+2p_2)$<br>$V_{\gamma} = \exp(1.5+4p_2)$ |

The core difference between *Precoss* and *Precoss-β* is the inversion scheme used for inference: Dynamic Expectation Maximisation (Friston, Trujillo-Barreto and Daunizeau, 2008) for Precoss, and Generalized filtering (Friston *et al.*, 2010) for *Precoss-β*. The latter features state-dependent precisions (Feldman and Friston, 2010), which we use to actively modulate the precision of bottom-up prediction errors of syllable and/or gamma units.

For details about common aspects for *Precoss* and *Precoss-β*, we refer to (Hovsepyan, Olasagasti and Giraud, 2020).

### Modulation of bottom-up information

To account for how modulation frequency of PEP affects the amount of non-redundant bottom-up information on syllable units, we defined a metric that captures when bottom-up evidence about the presence of a syllable is noteworthy. Specifically, we weigh the positive evidence in favor of a syllable by the level of “inactivity” of the corresponding syllable. If positive evidence about the presence of the syllable arrives when the corresponding syllable unit has low activity, it has a stronger contribution to the measure than when the same positive evidence arrives when the activity of the corresponding syllable unit is already high:

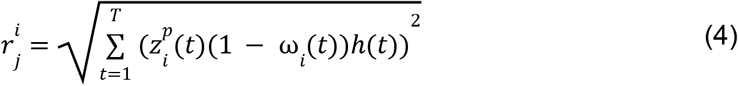

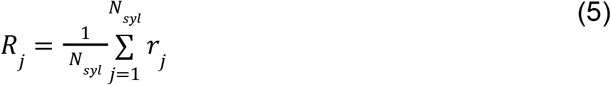

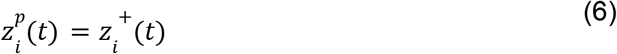

First, for each syllable *i* in sentence *j* we calculate the root sum square of the product of positive evidence in favor of syllable *i* (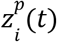, positive bottom-up prediction errors *z_i_*(*t*), equation 6) and its level of “inactivity” (1 – ω_*i*_(*t*)), where the summation is across time but excluding periods of active resetting of syllable units that happens when the gamma network signals the end of a syllable (for details see (Hovsepyan, Olasagasti and Giraud, 2020)) and ω_*i*_(*t*) corresponds to the *i-th* component of the softmax of the syllable hidden states (equation 4). *h*(*t*) is one outside active resetting periods and 0 during active resetting periods.

Then we define the average root sum square (*R_j_*) across all syllables (*N_syl_*) of sentence *j* (equation 5). *R_j_* is our measure of interest and was calculated for each Precoss-β variant and each modulation frequency.

### Statistical analysis

The model performance was evaluated based on the overlap measure (Supplementary Figure 1) that provides a single value for each sentence assessing the model’s ability to infer syllable identity and duration for each sentence. Simulations were performed on the same set of 220 sentences for each model variant and each frequency of modulation of prediction error precisions.

To compare the performance of *Precoss* vs *Precoss-β* we performed a Wilcoxon signed-rank test for each PEP frequency. To control for multiple comparisons the alpha = 0.05 was adjusted with the Bonferroni procedure. Each test was considered statistically significant if the p-value was less than 0.05/8 (dominator corresponds to the number of comparisons - the number of tested frequencies).

The same method was used for the comparisons reporting performance differences for comparing Precoss-β variants for each frequency of oscillating PEP (Supplementary Table 7) as well as for same-phase vs. anti-phase conditions (presented in Figure 5). Results are presented in Supplementary Tables 1-3 for *Precoss-β* variants A, B and C, and Supplementary Table 8 for anti-phase vs same-phase comparisons. In all tables the first column indicates which frequency is tested, the second column the associated signed-rank, and the third column the corresponding z-statistics. The last column represents the corresponding p-value.

For each *Precoss-β* variant, the effect of the modulation frequency was evaluated with a Friedman test, followed by multiple comparisons controlled by Bonferroni correction. Supplementary Tables 4-6 (for overlap measure) and 9-11 (non-redundant bottom-up information) report results of pairwise comparisons, where the first two columns indicate which modulation frequencies of precisions are being compared. The fourth column indicates the difference in the mean signed-rank for the corresponding pair, whereas the third and fifth columns indicate lower and upper bound of 95% confidence interval, correspondingly. Lastly, the sixth column represents Bonferroni corrected p-values. Pairwise comparisons are considered statistically significant if the corrected p-value < 0.05.

All statistical tests were performed using built-in Matlab functions. One sentence (N182) did not converge for *Precoss-β* (B), thus this sentence was excluded from all model variants in the calculation of BIC results (Figure 3) and the non-redundancy measure for variant B (Supplementary figure 7).

## Code and Data availability

Simulations were performed with DEM Toolbox in SPM (*SPM - Statistical Parametric Mapping,* no date) using MATLAB 2018b, The MathWorks, Inc., Natick, Massachusetts, United States. Custom code and raw data for statistical analysis would be freely distributed online.

## Acknowledgements

This work was funded by a grant from the NCCR Evolving Language, Swiss National Science Foundation Agreement #51NF40_180888. The computations were performed at University of Geneva on “Baobab” HPC cluster(s).

## Contributions

A.L.G. and I.O. designed the study, S.H. carried out the study, S.H., I.O., and A.L.G. wrote the manuscript.

