## Supplementary Information for "Rhythmic modulation of prediction errors: a possible role for the beta-range in speech processing"

#### Supplementary Figure 1

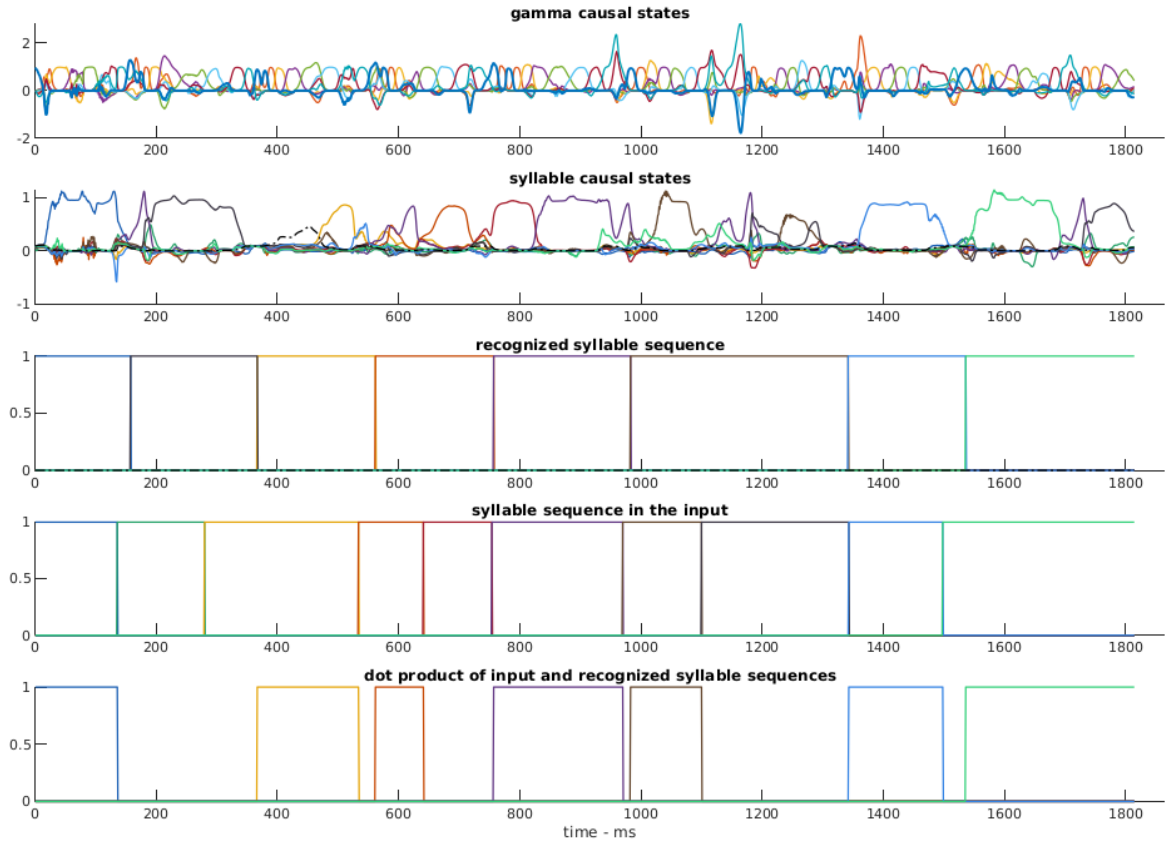


**Supplementary figure 1: Performance metric overlap based on the dynamics of syllable and gamma units** The top two panels represent the dynamics of the gamma and syllable units during the inference for an example sentence. For each subplot, coloured lines were used to represent different gamma and syllable units. The gamma unit with a thick blue line corresponds to the first gamma unit, whose peak (amplitude more than 0.6) is used as a marker to indicate windows for identifying the “winner” syllable unit. For the latter, we look for the syllable unit with the highest average activation within a gamma window (time interval between two consecutive gamma 1 peaks). The sequence of the recognized syllables is shown in the 3rd panel, whereas the sequence and duration of the syllables in the input are shown in the 4th subpanel. The model performance (the overlap metric) is evaluated with the dot-product (bottom subpanel) of recognized and input syllable sequences (subpanels 3 and 4) divided by the duration of the input sentence. The higher/closer to 1, the better the model is able to infer identity and duration of syllables in the input sentence.

### Supplementary Figure 2

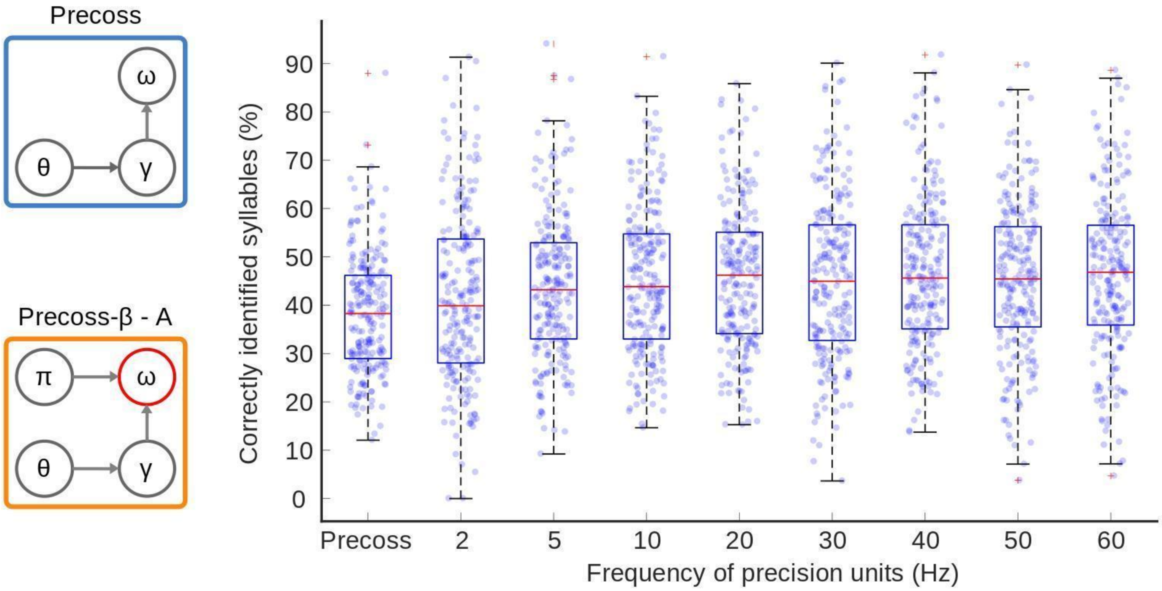


**Supplementary figure 2: *Precoss-β* (A) performance based on the overlap measure.** Simulation results on 220 sentences are presented. Performance is evaluated based on the overlap between the recognized syllable sequence and the sequence of syllables in the input sentence (for details, see Supplementary Figure 1). We compare the performance of the *Precoss-β* for different frequency values of PEP. For all frequencies, the model performance is better than the performance of *Precoss* with stationary precisions (Supplementary Table 1). Friedman test ($\chi 2$ = 22.89, $p$ = 0.0018) indicated an effect of PEP frequency on model performance. Post-hoc pairwise comparisons (Bonferroni-corrected, Supplementary Table 4), indicated that performance of *Precoss-β* increased with frequency up to 5 Hz and reached a plateau (there is no statistically significant difference in the model's performance for frequencies higher or equal to 5 Hz). Each point on the scatter plot represents the model performance in each sentence for the corresponding PEP frequency. The central-red mark of the box plots indicates the median, whereas bottom and top edges represent 25th and 75th percentiles. Red crosses indicate outliers, whereas whiskers extend to the highest and lowest overlap values that are not considered outliers.

### Supplementary Figure 3

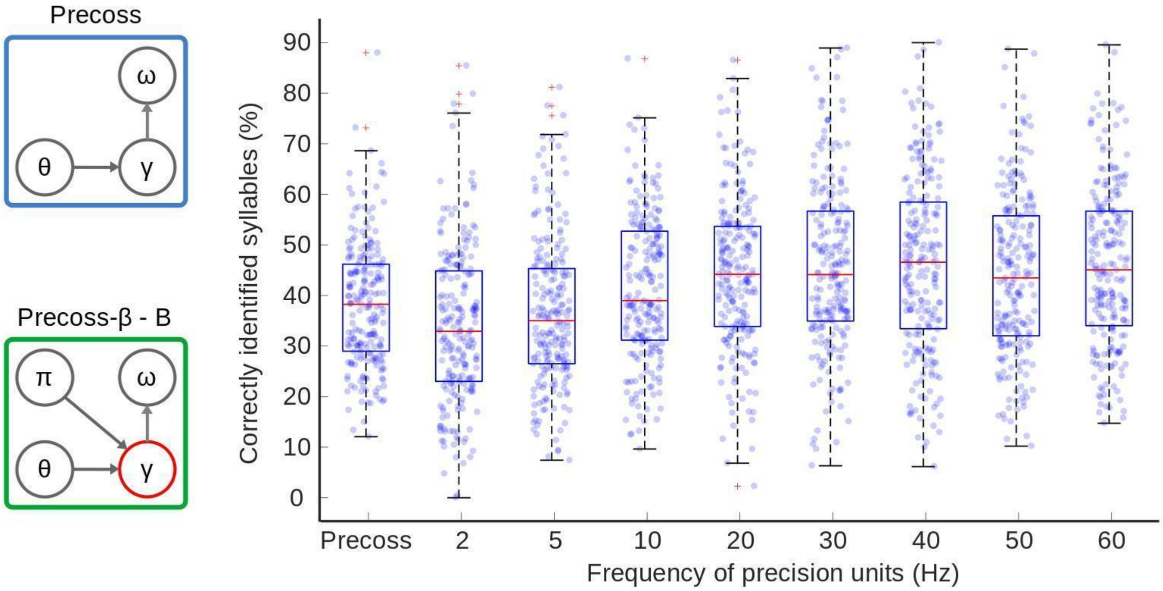


**Supplementary figure 3: *Precoss-β* (B) performance based on the overlap measure.** Simulation results on 220 sentences are presented in the figure. Performance is evaluated based on the overlap between the recognized and input syllable sequences (for details, see supplementary figure 1). *Precoss-β* outperforms *Precoss* for PEP frequencies higher or equal to 10 Hz, whereas for smaller frequencies the performance is worse (Supplementary Table 2). Friedman test ($\chi 2$ = 171.74, p = 1.076e-33) indicated an effect of PEP frequency on model performance. Post-hoc, multiple comparisons tests (corrected with Bonferroni procedure, Supplementary Table 5) indicated that the *Precoss-β* performance increases with the frequency and reaches a plateau at around 20 Hz. Each point on the scatter plot represents the measurement value for each sentence for the corresponding PEP frequency. The central-red mark of the box plots corresponds to the median, whereas bottom and top edges represent 25th and 75th percentiles, respectively. Red crosses indicate outliers, whereas whiskers extend to the highest and lowest performance values that are not considered outliers.

### Supplementary Figure 4

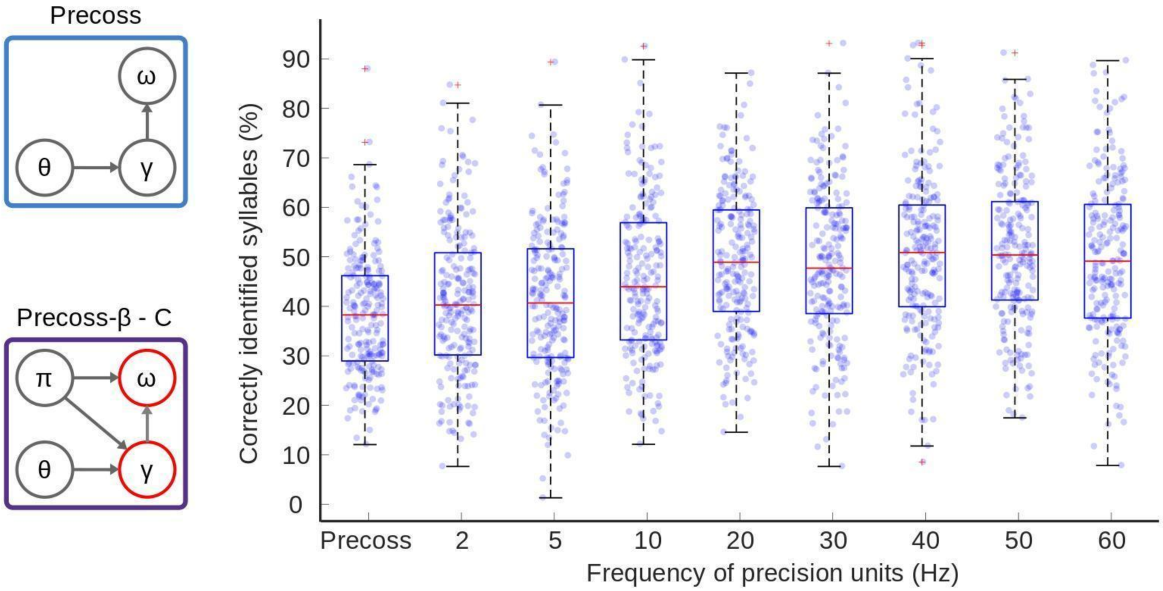


**Supplementary figure 4: *Precoss-β* (C) performance based on overlap measure.** Simulation results on 220 sentences are presented in the figure. Performance is evaluated based on the overlap duration between the recognized syllable sequence and the sequence of syllables in the input sentence (for details, see supplementary figure 1). For this condition performance of *Precoss-β* is better than the performance of Precoss, with stationary precisions for all frequency values of the precision units (Supplementary Table 3). Friedman test ($\chi2$ = 158.86, p = 5.577e-31) confirmed that the frequency of PEP affects model performance. Post-hoc, Bonferroni corrected pairwise comparisons indicated that the model performance increases with the frequency and reaches a plateau at 20 Hz (there are no statistically significant differences in performance for higher PEP frequencies, Supplementary Table 6). The central-red mark of the box plots corresponds to the median, whereas bottom and top edges represent 25th and 75th percentiles, respectively. Red crosses indicate outliers, whereas whiskers extend to the highest and lowest model performance values that are not considered outliers.

### Supplementary Figure 5

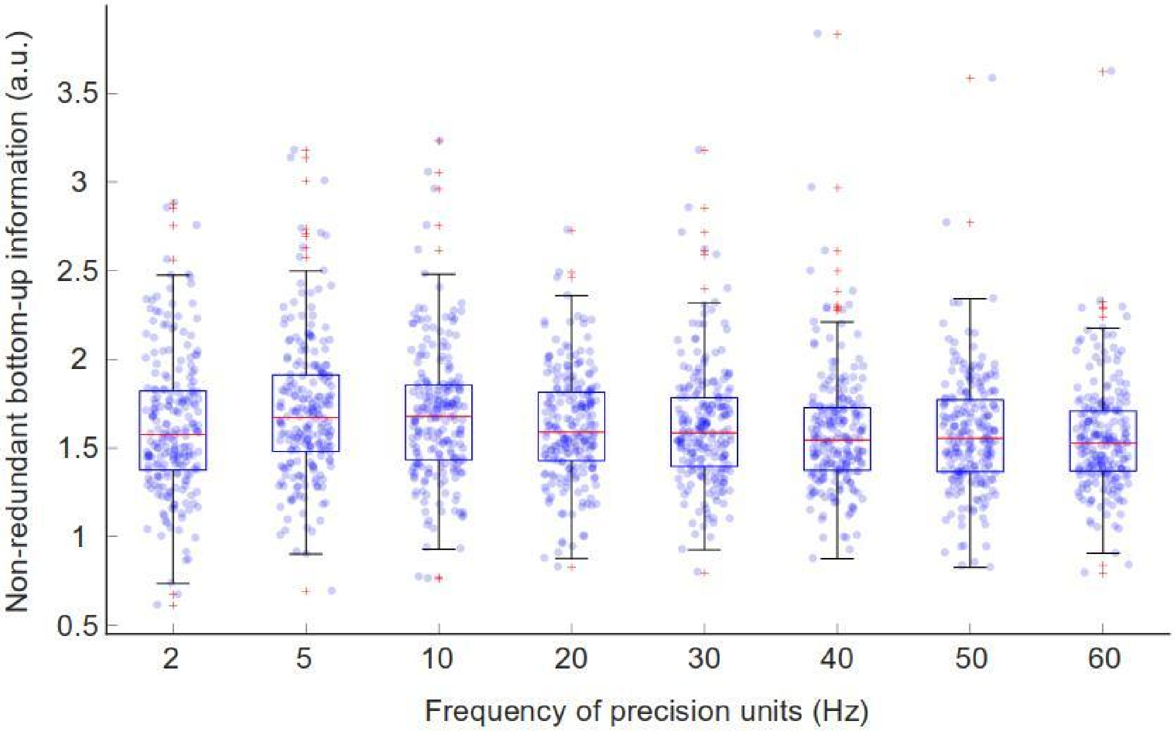


**Supplementary figure 5: Non-redundancy of bottom-up information - Precoss-β (A).** We evaluated how the modulation frequency of PEP affects the informativeness (non-redundancy) in the bottom-up prediction signal. We calculate the root sum square of bottom-up prediction errors while the corresponding syllable units are inactive (for details, see Methods). Friedman test indicated ($\chi^2$ = 65.18, p = 8.81e-14) that the modulation frequency affects the amount of informative signal propagated up in the model-hierarchy. Pairwise comparisons were performed for each frequency pair (Supplementary Table 9). Bonferroni procedure was used to control for multiple comparisons. The non-redundancy measure around 5-10 Hz was statistically higher ($p<0.05$) than for frequencies higher than 30 Hz. The central-red mark of the box plots corresponds to the median, whereas bottom and top edges represent 25th and 75th percentiles, respectively. Red crosses indicate outliers, whereas whiskers extend to the highest and lowest performance measures that are not considered outliers.

### Supplementary Figure 6

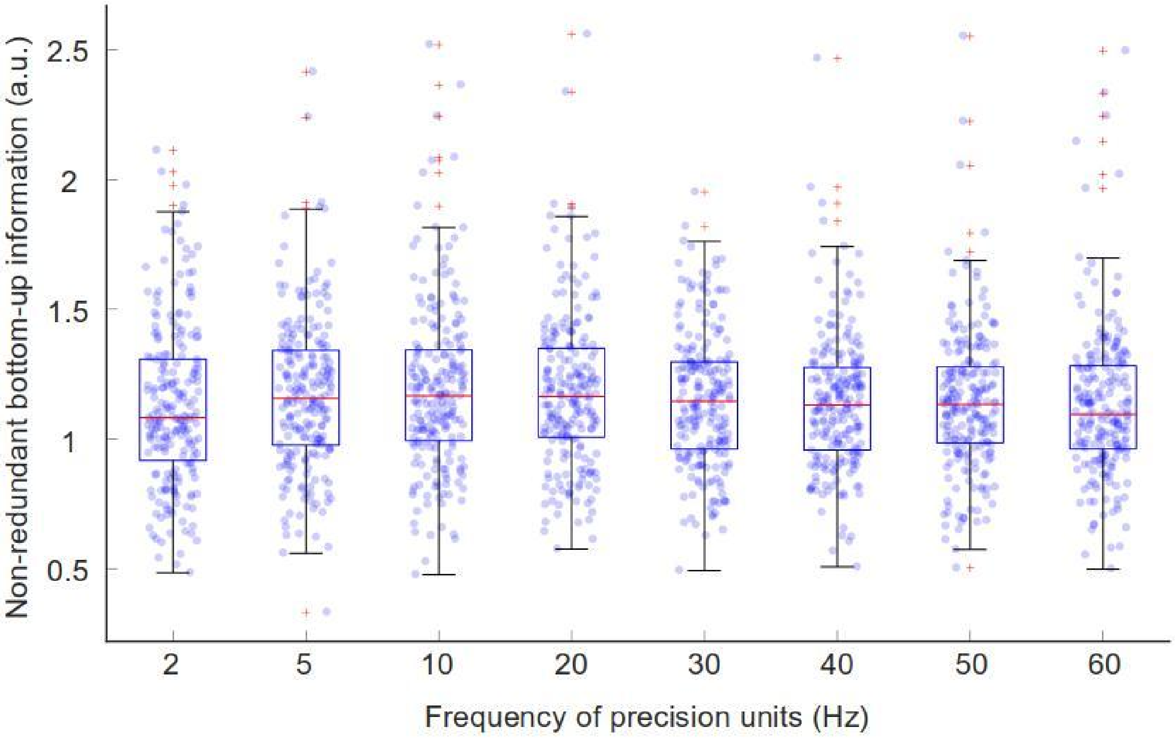


**Supplementary figure 6: Non-redundancy of bottom-up information - *Precoss-β* (B).** We evaluated how the modulation frequency of PEP affects the amount of informative (non-redundant) bottom-up prediction signal. We calculate the root sum square of bottom-up prediction errors while the corresponding syllable units are inactive (for details, see Methods). Friedman test indicated ($\chi^2 = 19.5$, $p = 0.0019$) that the modulation frequency affects the amount of informative signal propagated up in the model-hierarchy. Pairwise comparisons were performed for each frequency pair (Supplementary Table 10). Bonferroni procedure was used to control for multiple comparisons. Modulation frequencies 10, 20 Hz allowed for more bottom-up informative signals than the 5 Hz and 60 Hz ($p<0.05$). All other comparisons were statistically non-significant, as expected when precision is not modulated in syllable units. The central-red mark of the box plots corresponds to the median, whereas bottom and top edges represent 25th and 75th percentiles, respectively. Red crosses indicate outliers, whereas whiskers extend to the highest and lowest performance values that are not considered outliers.

#### Supplementary Table 1: *Precoss* vs. *Precoss-β* (A)

| frequency | signed rank | z-statistic | p-value |
|---|---|---|---|
| 2 Hz | 9293 | -2.8342 | 0.0046 |
| 5 Hz | 7561 | -4.7765 | 1.784e-6 |
| 10 Hz | 6256 | -6.2410 | 4.346e-10 |
| 20 Hz | 5926 | -6.5902 | 4.393e-11 |
| 30 Hz | 6524 | -5.9575 | 2.561e-9 |
| 40 Hz | 5596 | -6.9393 | 3.94e-12 |
| 50 Hz | 6502 | -5.9808 | 2.22e-9 |
| 60 Hz | 5888 | -6.6304 | 3.348e-11 |

Related to Figure 1.

#### Supplementary Table 2: *Precoss* vs. *Precoss-β* (B)

| frequency | signed rank | z-statistic | p-value |
|---|---|---|---|
| 2 Hz | 15342 | 3.6536 | 2.586e-4 |
| 5 Hz | 13316 | 1.2283 | 0.2193 |
| 10 Hz | 9286 | -2.9389 | 0.0033 |
| 20 Hz | 6907 | -5.5523 | 2.819e-8 |
| 30 Hz | 6568 | -5.9110 | 3.4e-9 |
| 40 Hz | 6192 | -6.3088 | 2.813e-10 |
| 50 Hz | 7565 | -4.8561 | 1.197e-6 |
| 60 Hz | 6033 | -6.4770 | 9.356e-11 |

Related to Figure 1.

Supplementary Table 3: *Precoss* vs. *Precoss-β* (C)

| frequency | signed rank | z-statistic | p-value |
|---|---|---|---|
| 2 Hz | 9469 | -2.7440 | 0.0061 |
| 5 Hz | 9481 | -2.8290 | 0.0047 |
| 10 Hz | 6272 | -6.2241 | 4.843e-10 |
| 20 Hz | 3448 | -9.2119 | 3.205e-20 |
| 30 Hz | 3991 | -8.6374 | 5.752e-18 |
| 40 Hz | 2554 | -10.11 | 4.987e-24 |
| 50 Hz | 2462 | -10.255 | 1.1234e-24 |
| 60 Hz | 3402 | -9.2605 | 2.034e-20 |

Related to Figure 1.

### Supplementary Table 4: Multiple comparison table for variant A - overlap measure

| freq.1 | freq. 2 | lower bound | mean diff. | upper bound | p-value |
|---|---|---|---|---|---|
| 2 Hz | 5 Hz | -1.149508141 | -0.4204545455 | 0.3085990502 | 1 |
| 2 Hz | 10 Hz | -1.526780868 | -0.7977272727 | -0.06867367707 | 1.77E-02 |
| 2 Hz | 20 Hz | -1.479053596 | -0.75 | -0.02094640434 | 3.67E-02 |
| 2 Hz | 30 Hz | -1.231326323 | -0.5022727273 | 0.2267808684 | 8.79E-01 |
| 2 Hz | 40 Hz | -1.669962687 | -0.9409090909 | -0.2118554953 | 1.55E-03 |
| 2 Hz | 50 Hz | -1.415417232 | -0.6863636364 | 0.04268995929 | 9.17E-02 |
| 2 Hz | 60 Hz | -1.522235414 | -0.7931818182 | -0.06412822253 | 1.90E-02 |
| 5 Hz | 10 Hz | -1.106326323 | -0.3772727273 | 0.3517808684 | 1 |
| 5 Hz | 20 Hz | -1.05859905 | -0.3295454545 | 0.3995081411 | 1.00E+00 |
| 5 Hz | 30 Hz | -0.8108717775 | -0.08181818182 | 0.6472354138 | 1.00E+00 |
| 5 Hz | 40 Hz | -1.249508141 | -0.5204545455 | 0.2085990502 | 7.21E-01 |
| 5 Hz | 50 Hz | -0.9949626866 | -0.2659090909 | 0.4631445047 | 1.00E+00 |
| 5 Hz | 60 Hz | -1.101780868 | -0.3727272727 | 0.3563263229 | 1.00E+00 |
| 10 Hz | 20 Hz | -0.6813263229 | 0.04772727273 | 0.7767808684 | 1 |
| 10 Hz | 30 Hz | -0.4335990502 | 0.2954545455 | 1.024508141 | 1 |
| 10 Hz | 40 Hz | -0.8722354138 | -0.1431818182 | 0.5858717775 | 1.00E+00 |
| 10 Hz | 50 Hz | -0.6176899593 | 0.1113636364 | 0.840417232 | 1.00E+00 |
| 10 Hz | 60 Hz | -0.7245081411 | 0.004545454545 | 0.7335990502 | 1 |
| 20 Hz | 30 Hz | -0.4813263229 | 0.2477272727 | 0.9767808684 | 1 |
| 20 Hz | 40 Hz | -0.9199626866 | -0.1909090909 | 0.5381445047 | 1 |
| 20 Hz | 50 Hz | -0.665417232 | 0.06363636364 | 0.7926899593 | 1 |
| 20 Hz | 60 Hz | -0.7722354138 | -0.04318181818 | 0.6858717775 | 1 |
| 30 Hz | 40 Hz | -1.167689959 | -0.4386363636 | 0.290417232 | 1 |
| 30 Hz | 50 Hz | -0.9131445047 | -0.1840909091 | 0.5449626866 | 1 |
| 30 Hz | 60 Hz | -1.019962687 | -0.2909090909 | 0.4381445047 | 1 |
| 40 Hz | 50 Hz | -0.4745081411 | 0.2545454545 | 0.9835990502 | 1 |
| 40 Hz | 60 Hz | -0.5813263229 | 0.1477272727 | 0.8767808684 | 1 |
| 50 Hz | 60 Hz | -0.8358717775 | -0.1068181818 | 0.6222354138 | 1 |

Related to Figure 1: Results of multiple comparisons followed the Friedman test.

### Supplementary Table 5: Multiple comparison table for variant B - overlap measure

| freq. 1 | freq. 2 | lower bound | mean diff. | upper bound | p-value |
|---|---|---|---|---|---|
| 2 Hz | 5 Hz | -1.222551381 | -0.4931818182 | 0.2361877442 | 0.9707576881 |
| 2 Hz | 10 Hz | -2.079369562 | -1.35 | -0.6206304377 | 2.07E-07 |
| 2 Hz | 20 Hz | -2.558915017 | -1.829545455 | -1.100175892 | 1.31E-13 |
| 2 Hz | 30 Hz | -2.833915017 | -2.104545455 | -1.375175892 | 5.60E-18 |
| 2 Hz | 40 Hz | -2.895278653 | -2.165909091 | -1.436539529 | 4.92E-19 |
| 2 Hz | 50 Hz | -2.504369562 | -1.775 | -1.045630438 | 8.17E-13 |
| 2 Hz | 60 Hz | -2.865733199 | -2.136363636 | -1.406994074 | 1.60E-18 |
| 5 Hz | 10 Hz | -1.586187744 | -0.8568181818 | -0.1274486195 | 0.006802847906 |
| 5 Hz | 20 Hz | -2.065733199 | -1.336363636 | -0.606994074 | 2.92E-07 |
| 5 Hz | 30 Hz | -2.340733199 | -1.611363636 | -0.881994074 | 1.44E-10 |
| 5 Hz | 40 Hz | -2.402096835 | -1.672727273 | -0.9433577104 | 2.20E-11 |
| 5 Hz | 50 Hz | -2.011187744 | -1.281818182 | -0.5524486195 | 1.13E-06 |
| 5 Hz | 60 Hz | -2.372551381 | -1.643181818 | -0.9138122558 | 5.48E-11 |
| 10 Hz | 20 Hz | -1.208915017 | -0.4795454545 | 0.2498241078 | 1 |
| 10 Hz | 30 Hz | -1.483915017 | -0.7545454545 | -0.0251758922 | 0.03447289972 |
| 10 Hz | 40 Hz | -1.545278653 | -0.8159090909 | -0.08653952857 | 1.33E-02 |
| 10 Hz | 50 Hz | -1.154369562 | -0.425 | 0.3043695623 | 1.00E+00 |
| 10 Hz | 60 Hz | -1.515733199 | -0.7863636364 | -0.05699407402 | 0.02121358755 |
| 20 Hz | 30 Hz | -1.004369562 | -0.275 | 0.4543695623 | 1 |
| 20 Hz | 40 Hz | -1.065733199 | -0.3363636364 | 0.393005926 | 1 |
| 20 Hz | 50 Hz | -0.6748241078 | 0.05454545455 | 0.7839150169 | 1 |
| 20 Hz | 60 Hz | -1.036187744 | -0.3068181818 | 0.4225513805 | 1 |
| 30 Hz | 40 Hz | -0.7907331987 | -0.06136363636 | 0.668005926 | 1 |
| 30 Hz | 50 Hz | -0.3998241078 | 0.3295454545 | 1.058915017 | 1 |
| 30 Hz | 60 Hz | -0.7611877442 | -0.03181818182 | 0.6975513805 | 1 |
| 40 Hz | 50 Hz | -0.3384604714 | 0.3909090909 | 1.120278653 | 1 |
| 40 Hz | 60 Hz | -0.6998241078 | 0.02954545455 | 0.7589150169 | 1 |
| 50 Hz | 60 Hz | -1.090733199 | -0.3613636364 | 0.368005926 | 1 |

Related to Figure 1: Results of multiple comparisons followed the Friedman test.

### Supplementary Table 6: Multiple comparison table for variant C - overlap measure

| freq. 1 | freq. 2 | lower bound | mean diff. | upper bound | p-value |
|---|---|---|---|---|---|
| 2 Hz | 5 Hz | -0.7589150169 | -0.02954545455 | 0.6998241078 | 1 |
| 2 Hz | 10 Hz | -1.536187744 | -0.8068181818 | -0.07744861948 | 0.01538360986 |
| 2 Hz | 20 Hz | -2.297551381 | -1.568181818 | -0.8388122558 | 5.22E-10 |
| 2 Hz | 30 Hz | -2.118005926 | -1.388636364 | -0.6592668013 | 7.64E-08 |
| 2 Hz | 40 Hz | -2.647551381 | -1.918181818 | -1.188812256 | 5.93E-15 |
| 2 Hz | 50 Hz | -2.688460471 | -1.959090909 | -1.229721347 | 1.36E-15 |
| 2 Hz | 60 Hz | -2.277096835 | -1.547727273 | -0.8183577104 | 9.49E-10 |
| 5 Hz | 10 Hz | -1.50664229 | -0.7772727273 | -0.04790316493 | 0.02441366992 |
| 5 Hz | 20 Hz | -2.268005926 | -1.538636364 | -0.8092668013 | 1.23E-09 |
| 5 Hz | 30 Hz | -2.088460471 | -1.359090909 | -0.6297213467 | 1.64E-07 |
| 5 Hz | 40 Hz | -2.618005926 | -1.888636364 | -1.159266801 | 1.69E-14 |
| 5 Hz | 50 Hz | -2.658915017 | -1.929545455 | -1.200175892 | 3.95E-15 |
| 5 Hz | 60 Hz | -2.247551381 | -1.518181818 | -0.7888122558 | 2.22E-09 |
| 10 Hz | 20 Hz | -1.490733199 | -0.7613636364 | -0.03199407402 | 0.03111210372 |
| 10 Hz | 30 Hz | -1.311187744 | -0.5818181818 | 0.1475513805 | 0.3558697591 |
| 10 Hz | 40 Hz | -1.840733199 | -1.111363636 | -0.381994074 | 5.43E-05 |
| 10 Hz | 50 Hz | -1.88164229 | -1.152272727 | -0.4229031649 | 2.24E-05 |
| 10 Hz | 60 Hz | -1.470278653 | -0.7409090909 | -0.01153952857 | 0.04222161737 |
| 20 Hz | 30 Hz | -0.5498241078 | 0.1795454545 | 0.9089150169 | 1 |
| 20 Hz | 40 Hz | -1.079369562 | -0.35 | 0.3793695623 | 1 |
| 20 Hz | 50 Hz | -1.120278653 | -0.3909090909 | 0.3384604714 | 1 |
| 20 Hz | 60 Hz | -0.7089150169 | 0.02045454545 | 0.7498241078 | 1 |
| 30 Hz | 40 Hz | -1.258915017 | -0.5295454545 | 0.1998241078 | 0.6533364083 |
| 30 Hz | 50 Hz | -1.299824108 | -0.5704545455 | 0.1589150169 | 0.4076876235 |
| 30 Hz | 60 Hz | -0.8884604714 | -0.1590909091 | 0.5702786533 | 1 |
| 40 Hz | 50 Hz | -0.7702786533 | -0.04090909091 | 0.6884604714 | 1 |
| 40 Hz | 60 Hz | -0.3589150169 | 0.3704545455 | 1.099824108 | 1 |
| 50 Hz | 60 Hz | -0.318005926 | 0.4113636364 | 1.140733199 | 1 |

Related to Figure 1: Results of multiple comparisons followed the Friedman test.

#### Supplementary Table 7: *Precoss-β* variants comparisons for each PEP frequency value

| frequency | variants | signed rank | z-statistic | p-value |
|---|---|---|---|---|
| 2 Hz | A vs B | 17517 | 5.6729 | 1.404e-8 |
| | C vs A | 11708 | -0.244 | 0.8072 |
| | C vs B | 17474 | 5.9403 | 2.8454e-9 |
| 5 Hz | A vs B | 17915 | 6.094 | 1.1013e-9 |
| | C vs A | 10020 | -2.2588 | 0.0239 |
| | C vs B | 15970 | 4.0362 | 5.4322e-5 |
| 10 Hz | A vs B | 14980 | 2.9888 | 0.0028 |
| | C vs A | 12394 | 0.3718 | 0.7101 |
| | C vs B | 14896 | 3.0369 | 0.0024 |
| 20 Hz | A vs B | 12812 | 0.817 | 0.4139 |
| | C vs A | 15193 | 3.2142 | 0.0013 |
| | C vs B | 15774 | 3.8288 | 1.2875e-4 |
| 30 Hz | A vs B | 11659 | -0.5248 | 0.5998 |
| | C vs A | 14644 | 2.905 | 0.0037 |
| | C vs B | 14508 | 2.4894 | 0.0128 |
| 40 Hz | A vs B | 12005 | -0.0426 | 0.966 |
| | C vs A | 15511 | 3.5506 | 3.8437e-4 |
| | C vs B | 15515 | 3.5543 | 3.7899e-4 |
| 50 Hz | A vs B | 12005 | 0.9182 | 0.3585 |
| | C vs A | 16796 | 4.9101 | 9.103e-7 |
| | C vs B | 17794 | 5.966 | 2.4319e-9 |
| 60 Hz | A vs B | 12341 | 0.1968 | 0.844 |
| | C vs A | 15325 | 3.2586 | 0.0011 |
| | C vs B | 15928 | 3.9918 | 6.5582e-5 |

Related to Figure 1. For each frequency value, the performance difference is considered statistically significant if the $p < 0.05/3$ (corrected for multiple comparisons with Bonferroni procedure)

### Supplementary Table 8: Effect of the oscillating PEP phase on model performance

| frequency | signed rank | z-statistic | p-value |
|---|---|---|---|
| 2 Hz | 6576 | -5.9025 | 3.5806e-9 |
| 5 Hz | 9062 | -3.2723 | 0.0011 |
| 10 Hz | 8441 | -3.9294 | 0.0001 |
| 20 Hz | 9175 | -3.1528 | 0.0016 |
| 30 Hz | 9502 | -2.8068 | 0.0050 |
| 40 Hz | 7255 | -5.1024 | 3.3535e-7 |
| 50 Hz | 7140 | -5.3058 | 1.1219e-7 |
| 60 Hz | 7672 | -4.7429 | 2.1064e-6 |

Related to Figure 5. The performance difference is considered statistically significant if the $p < 0.05/8$ (corrected for multiple comparisons with Bonferroni procedure)

#### Supplementary Table 9: Multiple comparison table for variant A - (non-redundancy of bottom-up information)

| freq. 1 | freq. 2 | lower bound | mean diff. | upper bound | p-value |
|---|---|---|---|---|---|
| 2 Hz | 5 Hz | -1.325001779 | -0.5954545455 | 0.134092688 | 0.3019842359 |
| 2 Hz | 10 Hz | -1.306819961 | -0.5772727273 | 0.1522745062 | 3.76E-01 |
| 2 Hz | 20 Hz | -0.7431835971 | -0.01363636364 | 0.7159108698 | 1.00E+00 |
| 2 Hz | 30 Hz | -0.6250017789 | 0.1045454545 | 0.834092688 | 1.00E+00 |
| 2 Hz | 40 Hz | -0.2022745062 | 0.5272727273 | 1.256819961 | 6.71E-01 |
| 2 Hz | 50 Hz | -0.09772905165 | 0.6318181818 | 1.361365415 | 1.91E-01 |
| 2 Hz | 60 Hz | 0.1749982211 | 0.9045454545 | 1.634092688 | 3.01E-03 |
| 5 Hz | 10 Hz | -0.7113654153 | 0.01818181818 | 0.7477290516 | 1 |
| 5 Hz | 20 Hz | -0.1477290516 | 0.5818181818 | 1.311365415 | 3.56E-01 |
| 5 Hz | 30 Hz | -0.02954723346 | 0.7 | 1.429547233 | 7.63E-02 |
| 5 Hz | 40 Hz | 0.3931800393 | 1.122727273 | 1.852274506 | 4.28E-05 |
| 5 Hz | 50 Hz | 0.4977254938 | 1.227272727 | 1.956819961 | 4.15E-06 |
| 5 Hz | 60 Hz | 0.7704527665 | 1.5 | 2.229547233 | 3.75E-09 |
| 10 Hz | 20 Hz | -0.1659108698 | 0.5636363636 | 1.293183597 | 0.442589689 |
| 10 Hz | 30 Hz | -0.04772905165 | 0.6818181818 | 1.411365415 | 0.09820677438 |
| 10 Hz | 40 Hz | 0.3749982211 | 1.104545455 | 1.834092688 | 6.31E-05 |
| 10 Hz | 50 Hz | 0.4795436756 | 1.209090909 | 1.938638143 | 6.31E-06 |
| 10 Hz | 60 Hz | 0.7522709484 | 1.481818182 | 2.211365415 | 6.24E-09 |
| 20 Hz | 30 Hz | -0.6113654153 | 0.1181818182 | 0.8477290516 | 1 |
| 20 Hz | 40 Hz | -0.1886381426 | 0.5409090909 | 1.270456324 | 0.5755789975 |
| 20 Hz | 50 Hz | -0.08409268801 | 0.6454545455 | 1.375001779 | 0.1600342562 |
| 20 Hz | 60 Hz | 0.1886345847 | 0.9181818182 | 1.647729052 | 0.002364479987 |
| 30 Hz | 40 Hz | -0.3068199607 | 0.4227272727 | 1.152274506 | 1 |
| 30 Hz | 50 Hz | -0.2022745062 | 0.5272727273 | 1.256819961 | 0.671095035 |
| 30 Hz | 60 Hz | 0.07045276654 | 0.8 | 1.529547233 | 0.0171893505 |
| 40 Hz | 50 Hz | -0.6250017789 | 0.1045454545 | 0.834092688 | 1 |
| 40 Hz | 60 Hz | -0.3522745062 | 0.3772727273 | 1.106819961 | 1 |
| 50 Hz | 60 Hz | -0.4568199607 | 0.2727272727 | 1.002274506 | 1 |

Related to Figure 4.

#### Supplementary Table 10: Multiple comparison table for variant B - (non-redundancy of bottom-up information)

| freq. 1 | freq. 2 | lower bound | mean diff. | upper bound | p-value |
|---|---|---|---|---|---|
| 2 Hz | 5 Hz | -1.178699554 | -0.4474885845 | 0.2837223849 | 1 |
| 2 Hz | 10 Hz | -1.466370787 | -0.7351598174 | -0.003948848013 | 4.72E-02 |
| 2 Hz | 20 Hz | -1.493768047 | -0.7625570776 | -0.03134610829 | 3.15E-02 |
| 2 Hz | 30 Hz | -1.151302294 | -0.4200913242 | 0.3111196451 | 1.00E+00 |
| 2 Hz | 40 Hz | -1.064544303 | -0.3333333333 | 0.397877636 | 1.00E+00 |
| 2 Hz | 50 Hz | -0.9321242113 | -0.200913242 | 0.5302977273 | 1.00E+00 |
| 2 Hz | 60 Hz | -0.699247499 | 0.03196347032 | 0.7631744397 | 1.00E+00 |
| 5 Hz | 10 Hz | -1.018882202 | -0.2876712329 | 0.4435397365 | 1 |
| 5 Hz | 20 Hz | -1.046279462 | -0.3150684932 | 0.4161424762 | 1.00E+00 |
| 5 Hz | 30 Hz | -0.7038137091 | 0.02739726027 | 0.7586082296 | 1.00E+00 |
| 5 Hz | 40 Hz | -0.6170557182 | 0.1141552511 | 0.8453662205 | 1.00E+00 |
| 5 Hz | 50 Hz | -0.4846356269 | 0.2465753425 | 0.9777863118 | 1.00E+00 |
| 5 Hz | 60 Hz | -0.2517589145 | 0.4794520548 | 1.210663024 | 1.00E+00 |
| 10 Hz | 20 Hz | -0.7586082296 | -0.02739726027 | 0.7038137091 | 1 |
| 10 Hz | 30 Hz | -0.4161424762 | 0.3150684932 | 1.046279462 | 1 |
| 10 Hz | 40 Hz | -0.3293844853 | 0.401826484 | 1.133037453 | 1.00E+00 |
| 10 Hz | 50 Hz | -0.196964394 | 0.5342465753 | 1.265457545 | 6.29E-01 |
| 10 Hz | 60 Hz | 0.03591231833 | 0.7671232877 | 1.498334257 | 2.94E-02 |
| 20 Hz | 30 Hz | -0.3887452159 | 0.3424657534 | 1.073676723 | 1 |
| 20 Hz | 40 Hz | -0.301987225 | 0.4292237443 | 1.160434714 | 1 |
| 20 Hz | 50 Hz | -0.1695671337 | 0.5616438356 | 1.292854805 | 0.4598859512 |
| 20 Hz | 60 Hz | 0.06330957861 | 0.7945205479 | 1.525731517 | 0.01927270741 |
| 30 Hz | 40 Hz | -0.6444529785 | 0.08675799087 | 0.8179689602 | 1 |
| 30 Hz | 50 Hz | -0.5120328871 | 0.2191780822 | 0.9503890515 | 1 |
| 30 Hz | 60 Hz | -0.2791561748 | 0.4520547945 | 1.183265764 | 1 |
| 40 Hz | 50 Hz | -0.598790878 | 0.1324200913 | 0.8636310607 | 1 |
| 40 Hz | 60 Hz | -0.3659141657 | 0.3652968037 | 1.096507773 | 1 |
| 50 Hz | 60 Hz | -0.498334257 | 0.2328767123 | 0.9640876817 | 1 |

Related to Supplementary Figure 5.

#### Supplementary Table 11: Multiple comparison table for variant C - (non-redundancy of bottom-up information)

| freq. 1 | freq. 2 | lower bound | mean diff. | upper bound | p-value |
|---|---|---|---|---|---|
| 2 Hz | 5 Hz | -2.306819961 | -1.577272727 | -0.8477254938 | 4.04E-10 |
| 2 Hz | 10 Hz | -3.106819961 | -2.377272727 | -1.647725494 | 6.90E-23 |
| 2 Hz | 20 Hz | -3.925001779 | -3.195454545 | -2.465907312 | 3.64E-41 |
| 2 Hz | 30 Hz | -3.875001779 | -3.145454545 | -2.415907312 | 6.75E-40 |
| 2 Hz | 40 Hz | -3.211365415 | -2.481818182 | -1.752270948 | 6.28E-25 |
| 2 Hz | 50 Hz | -2.625001779 | -1.895454545 | -1.165907312 | 1.35E-14 |
| 2 Hz | 60 Hz | -2.456819961 | -1.727272727 | -0.9977254938 | 3.94E-12 |
| 5 Hz | 10 Hz | -1.529547233 | -0.8 | -0.07045276654 | 0.0171893505 |
| 5 Hz | 20 Hz | -2.347729052 | -1.618181818 | -0.8886345847 | 1.19E-10 |
| 5 Hz | 30 Hz | -2.297729052 | -1.568181818 | -0.8386345847 | 5.28E-10 |
| 5 Hz | 40 Hz | -1.634092688 | -0.9045454545 | -0.1749982211 | 3.01E-03 |
| 5 Hz | 50 Hz | -1.047729052 | -0.3181818182 | 0.4113654153 | 1.00E+00 |
| 5 Hz | 60 Hz | -0.8795472335 | -0.15 | 0.5795472335 | 1.00E+00 |
| 10 Hz | 20 Hz | -1.547729052 | -0.8181818182 | -0.08863458472 | 0.01286952994 |
| 10 Hz | 30 Hz | -1.497729052 | -0.7681818182 | -0.03863458472 | 0.02813652307 |
| 10 Hz | 40 Hz | -0.834092688 | -0.1045454545 | 0.6250017789 | 1.00E+00 |
| 10 Hz | 50 Hz | -0.2477290516 | 0.4818181818 | 1.211365415 | 1.00E+00 |
| 10 Hz | 60 Hz | -0.07954723346 | 0.65 | 1.379547233 | 1.51E-01 |
| 20 Hz | 30 Hz | -0.6795472335 | 0.05 | 0.7795472335 | 1 |
| 20 Hz | 40 Hz | -0.01591086983 | 0.7136363636 | 1.443183597 | 0.06288924011 |
| 20 Hz | 50 Hz | 0.5704527665 | 1.3 | 2.029547233 | 7.29E-07 |
| 20 Hz | 60 Hz | 0.7386345847 | 1.468181818 | 2.197729052 | 9.10E-09 |
| 30 Hz | 40 Hz | -0.06591086983 | 0.6636363636 | 1.393183597 | 0.1257168479 |
| 30 Hz | 50 Hz | 0.5204527665 | 1.25 | 1.979547233 | 2.43E-06 |
| 30 Hz | 60 Hz | 0.6886345847 | 1.418181818 | 2.147729052 | 3.53E-08 |
| 40 Hz | 50 Hz | -0.1431835971 | 0.5863636364 | 1.31591087 | 0.3374168216 |
| 40 Hz | 60 Hz | 0.02499822108 | 0.7545454545 | 1.484092688 | 0.0345679525 |
| 50 Hz | 60 Hz | -0.5613654153 | 0.1681818182 | 0.8977290516 | 1 |

Related to Supplementary Figure 6.
